# An AlphaFold2 map of the 53BP1 pathway identifies a direct SHLD3-RIF1 interaction critical for DNA repair activity

**DOI:** 10.1101/2023.01.12.523815

**Authors:** Chérine Sifri, Lisa Hoeg, Daniel Durocher, Dheva Setiaputra

**Affiliations:** Lunenfeld-Tanenbaum Research Institute, Mount Sinai Hospital, 600 University Avenue, Toronto, ON, M5G 1X5, Canada; Department of Biochemistry, University of Toronto, 1 King’s College Circle, Toronto, ON, M5S 1A8, Canada; Department of Molecular Genetics, University of Toronto, 1 King’s College Circle, Toronto, ON, M5S 1A8, Canada

**Author notes:** Co-corresponding authors (,), Lead Contact: Daniel Durocher, Format: Scientific Report.

## Abstract

53BP1 is a chromatin-binding DNA repair protein that promotes DNA double-strand break repair through recruitment of downstream effectors including RIF1, shieldin, and CST. The structural basis of the protein-protein interactions within the 53BP1-RIF1-shieldin-CST pathway that are essential for its DNA repair activity are largely unknown. Here we used AlphaFold2-Multimer (AF2) to predict all possible pairwise combinations of proteins within this pathway and provide structural models of seven previously characterized interactions. This analysis also predicted an entirely novel binding interface between the HEAT-repeat domain of RIF1 and the eIF4E-like domain of SHLD3. Extensive interrogation of this interface through both in vitro pulldown analysis and cellular assays supports the AF2-predicted model and demonstrates that RIF1-SHLD3 binding is essential for shieldin recruitment to sites of DNA damage, and for its role in antibody class switch recombination. Direct physical interaction between RIF1 and SHLD3 is therefore essential for 53BP1-RIF1-shieldin-CST pathway activity.

## Introduction

The nucleolytic processing and fill-in synthesis of the ends of DNA double-strand breaks (DSBs) is under the control of the chromatin-binding protein 53BP1, which is recruited to RNF168-ubiquitylated chromatin in a large domain that flanks the DSB (Panier and Boulton, 2014). The regulation of DNA end processing (or stability) impacts the type of DSB repair employed to mend the break, with 53BP1 promoting DNA end-stability which in turn favors non-homologous end-joining (NHEJ). 53BP1 carries out this function by acting as a platform for the recruitment of downstream factors that include RIF1 and PTIP (Callen et al., 2013; Chapman et al., 2013; Escribano-Díaz et al., 2013; Zimmermann et al., 2013). RIF1 then recruits the shieldin complex to facilitate DNA repair (Dev et al., 2018; Gao et al., 2018; Ghezraoui et al., 2018; Gupta et al., 2018; Mirman et al., 2018; Noordermeer et al., 2018; Setiaputra and Durocher, 2019). The 53BP1-RIF1-shieldin pathway promotes NHEJ and DSB end-stability by opposing the 5’-3’ nuclease degradation of DSB ends associated with homologous recombination (HR) repair. 53BP1-RIF1-shieldin-stimulated NHEJ is essential for antibody class switch recombination in B cells (CSR; Chapman et al., 2013; Ghezraoui et al., 2018; Noordermeer et al., 2018; Ward et al., 2004). Its role in suppressing HR mediates the toxicity of poly(ADP) ribose polymerase 1 inhibitors (PARPi) in *BRCA1*-mutated tumors (Noordermeer et al., 2018). Furthermore, 53BP1 accumulates in large nuclear bodies in G1 following DNA replication stress during the preceding cell cycle and suppresses toxic HR-mediated repair at these lesions (Spies et al., 2019). Although the precise mechanism underlying HR suppression by this pathway remains under investigation, shieldin recruits the CTC1-STN1-TEN1 (CST)-Pol α complex to DSBs to perform fill-in DNA synthesis at resected breaks (Mirman et al., 2018, 2022b; Schimmel et al., 2021). Shieldin has also been shown to recruit the structure-specific endonuclease ASTE1 to DSBs where it cleaves single-stranded DNA (ssDNA) overhangs (Zhao et al., 2021). Understanding the molecular basis underlying the recruitment of 53BP1 and its downstream effectors to DSBs is essential to determine how this pathway regulates DNA repair.

53BP1 is recruited to DNA damage sites by directly binding methylated histone H4 and DNA damage-induced histone H2A lysine 15 ubiquitylation through its tandem Tudor and ubiquitin-dependent recruitment domains, respectively (Fradet-Turcotte et al., 2013). RIF1 localizes to DNA breaks by binding three doubly-phosphorylated 53BP1 epitopes that are characterized by an LxL dileucine motif (x signifies any residue) preceding the phosphoresidues (Setiaputra et al., 2022). RIF1 recognizes phosphorylated 53BP1 using its N-terminal HEAT repeat domain (Setiaputra et al., 2022). Precisely how shieldin is recruited to DSBs is unknown, although RIF1 and shieldin can interact (Noordermeer et al., 2018; Setiaputra et al., 2022), suggesting that a hitherto uncharacterized RIF1-shieldin binding interface mediates shieldin localization to and function at DNA break sites. Shieldin consists of four subunits: SHLD1, SHLD2, SHLD3, and REV7 (Setiaputra and Durocher, 2019). SHLD3 is required for DSB localization of all other shieldin subunits, making it the primary candidate for RIF1-mediated recruitment (Noordermeer et al., 2018). SHLD3 binds REV7 through its N-terminus, and REV7 in turn binds the SHLD2 N-terminus (Dai et al., 2020; Ghezraoui et al., 2018; Liang et al., 2020). The SHLD2 C-terminus consists of three tandem oligonucleotide-binding folds (OB-folds) that binds ssDNA and serves as the binding site of SHLD1 (Noordermeer et al., 2018). Yeast two-hybrid experiments found that CST binds shieldin at multiple sites, with SHLD1 providing a key interaction interface (Mirman et al., 2018, 2022b). Aside from the nucleosome-53BP1 and SHLD3-REV7-SHLD2 subcomplexes (Liang et al., 2020; Wilson et al., 2016), the structural basis for the protein-protein interactions outlined above are unknown.

In this study, we used the protein-protein interaction prediction algorithm AlphaFold2-Multimer (AF2; Evans et al., 2021; Jumper et al., 2021; Mirdita et al., 2022) to probe the 53BP1-RIF1-shieldin-CST pathway. This approach accurately predicts known binding interfaces and provides the structural basis for multiple previously characterized interactions within the 53BP1 pathway. Additionally, AF2 analysis predicted an entirely novel interaction between the eIF4E-like domain of SHLD3 and the HEAT-repeat domain of RIF1. We confirmed this prediction both *in vitro* and in cellular assays measuring shieldin recruitment to sites of DNA damage by RIF1 and show that these mutations also disrupt shieldin-dependent CSR. Shieldin recruitment to DSB sites, and its ability to modulate DSB repair is therefore dependent on a direct SHLD3-RIF1 interaction.

## Results and Discussion

### Modeling the 53BP1-RIF1-shieldin-CST interaction network using AlphaFold2

We sought to understand the structural basis for the interactions facilitating 53BP1-RIF1-shieldin-CST recruitment to sites of DNA damage. Despite a good understanding of the domains involved in these interactions, the large size and propensity for disorder within the proteins of this pathway makes them refractory for purification and structural determination. We instead used AlphaFold2-Multimer to predict the heterodimeric structure for every unique pairwise combination for all known components of this pathway (53BP1, RIF1, SHLD1, SHLD2, SHLD3, REV7, ASTE1, CTC1, STN1, TEN1) (Fig 1A). Due to graphical memory limitations, we divided large proteins into multiple fragments. Five models were generated for each prediction and we scored each model based on pDockQ and mean interface predicted aligned error (PAE), two parameters that discriminate against incorrect AF2 predictions (Fig S1A; Bryant et al., 2022; Yin et al., 2022). Out of 855 models from 171 unique protein pairs, 95 models (11.1%) from 31 pairs satisfied the pDockQ and mean interface PAE cutoffs of >0.23 and <15 Å, respectively (Fig S1B). To further remove false positives, we assigned higher prediction confidence to interaction pairs in which ≥4 out of 5 models were consistent with each other. Predictions meeting this cutoff are depicted in Fig 1B, with the addition of the 53BP1 oligomerization domain which only has 3/5 consistent models but is supported by experimental evidence (Sundaravinayagam et al., 2019; Zgheib et al., 2009). Where possible, we merged multiple complementary pairwise predictions to depict larger macromolecular complexes (SHLD1-SHLD2-CTC1, CTC1-STN1-TEN1; Fig 1B). This analysis successfully recapitulated experimentally determined structures of SHLD3-REV7 (PDB ID:6KTO; Liang et al., 2020) and the CST complex consisting of CTC1-STN1-TEN1 (PDB ID:6W6W; Lim et al., 2020), further supporting the utility of this approach. Interestingly, the AF2 analysis predicted multiple unusual REV7 structures. REV7 is known to form head-to-head dimers (Xie et al., 2021), but AF2 predicted an extensively domain-swapped dimer instead (Fig S1C). Furthermore, REV7 is predicted to interact with four unique RIF1 and 53BP1 fragments, and all four predict the same mode of HORMA domain seatbelt-mediated interaction seen in REV7-SHLD3 (Fig S1D; Liang et al., 2020). Since we previously showed that REV7 requires SHLD3 for 53BP1 or RIF1 colocalization at DSBs (Noordermeer et al., 2018), we excluded these models from further analysis. ASTE1 was not predicted to interact with any proteins tested (Fig S1B).

**Figure 1.**
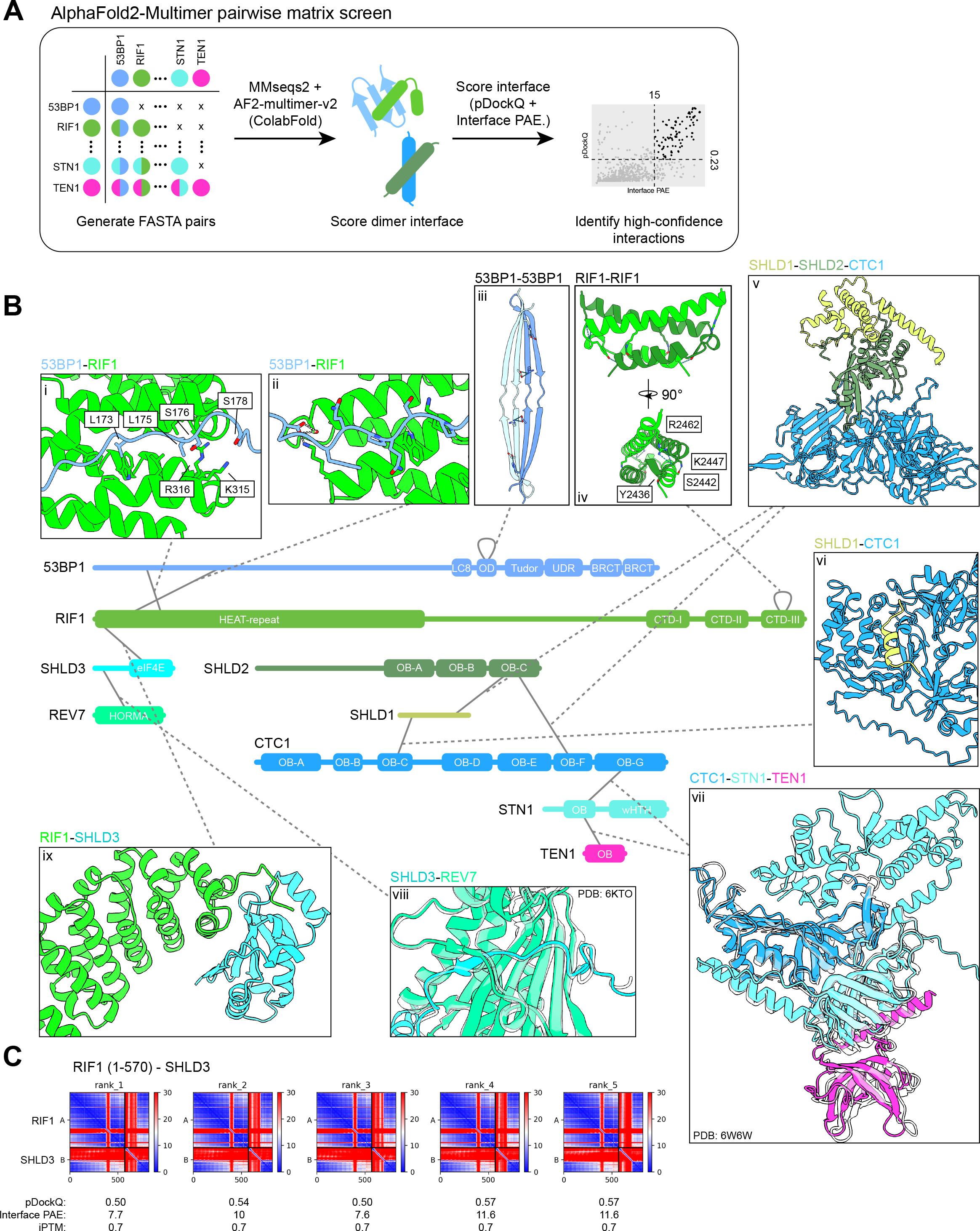
Exhaustive AlphaFold2-Multimer prediction of pairwise protein-protein interactions within the 53BP1-RIF1-shieldin-CST pathway. A. Schematic of the AF2 pairwise matrix screen for protein-protein interactions within the 53BP1-RIF1-shieldin-CST pathway. B. High-confidence interactions predicted by AF2. Structures of interfaces are shown, with corresponding experimental structures overlaid (translucent) if available (CST complex and SHLD3-REV7; PDB ID: 6W6W, 6KTO). Modeled hydrogen bonds are shown as dashed lines. See also Fig S1. C. Predicted aligned error (PAE) plots of RIF1 (1-570) and SHLD3, with calculated pDockQ, interface PAE, and interface predicted template modeling (iPTM) scores.

### Structural basis for 53BP1-RIF1 interaction and oligomerization

53BP1 is predicted to interact with RIF1 through its N-terminal unstructured domain and with itself through its oligomerization domain (Fig 1B). We previously identified that RIF1 binds three 53BP1 phosphorylated epitopes that mediate its recruitment to DSB sites (Setiaputra et al., 2022). Strikingly, despite not being designed to identify post-translational modifications, AF2 identified two of these motifs (Fig 1B panel i, S1E-H). In both cases, the apposed serine and leucine residues are oriented towards a groove formed between two RIF1 α-helical HEAT repeats (Fig S1G). Remarkably, the two 53BP1 phosphoacceptor serine residues are oriented towards the RIF1 K315/R316 residues that we found to be essential for this interaction (Setiaputra et al., 2022). The predicted structures comfortably accommodate phosphate groups modeled onto the putatively phosphorylated serines (Fig S1G, left panel). The two leucine residues of the LxL motif, which are essential for RIF1-53BP1 binding, are buried within a RIF1 hydrophobic cleft (Fig S1G, right panel). Interestingly, all five AF2 models predict a second site of 53BP1-RIF1 interaction involving 53BP1 residues 440-462 (Fig 1B panel ii, S1E-F). This region contains one predicted ATM phosphosite at S452. This residue is neither essential nor sufficient for RIF1 recruitment to DNA damage sites (Setiaputra et al., 2022), and could potentially represent a secondary 53BP1-RIF1 interaction site. These AF2 predictions provide a compelling structural explanation for the RIF1-53BP1 phosphodependent interaction that we previously described.

53BP1 and RIF1 are both known to form higher ordered structures (Moriyama et al., 2018; Zgheib et al., 2009). 53BP1 multimerization through its oligomerization domain (OD) is important for its DNA damage localization and function (Sundaravinayagam et al., 2019; Zgheib et al., 2009). The AF2 prediction corresponding to this region predicted that the 53BP1-OD dimerizes through an extended antiparallel beta sheet (Fig 1B panel iii) stabilized through backbone interactions and two hydrogen bonds between highly conserved residues (D1256-H1239 and E1245-R1252; Fig S1I). Notably, D1256 is essential for 53BP1 oligomerization and DNA damage recruitment, supporting this prediction (Zgheib et al., 2009). The 53BP1-OD tetramerizes *in vitro* (Sundaravinayagam et al., 2019), but subsequent AF2 prediction using four copies of the domain did not recapitulate a tetrameric arrangement. Nevertheless, the predicted 53BP1-OD dimer potentially represents a sub-assembly within the tetrameric structure.

RIF1 is known to dimerize through both its N-terminal HEAT repeats and its extreme C-terminus (Moriyama et al., 2018). The AF2 analysis predicted C-terminus dimerization through a head-to-head intercalated four-helix bundle stabilized by a buried hydrophobic core and two hydrogen bonds (Fig 1B panel iv, S1J). These interactions involve highly conserved hydrophobic and polar residues, respectively (L2417, I2421, L2424, L2441, L2448, V2455 and S2422, Y2436, R2462, K2447; Fig S1K). This region is conserved in yeast Rif1, which oligomerizes in an analogous fashion (Fig S1L; Shi et al., 2013). The biological importance of RIF1 dimerization is unclear and identifying specific residues necessary for this interaction will be essential for future efforts to address this question.

### Predicted interactions between the shieldin and CST complexes

The molecular determinants of shieldin and CST complex assembly are relatively well understood. The AF2 analysis recapitulated the interaction between the SHLD3 N-terminus and the REV7 HORMA domain (Liang et al., 2020), though it did not detect the interaction between the SHLD2 N-terminus and REV7. SHLD1 is known to interact with the SHLD2 C-terminus which contains three tandem OB-folds, specifically through the third OB-fold (Dev et al., 2018). Consistent with this experimental observation, AF2 predicts that SHLD1 binds the SHLD2 third OB-fold (Fig 1B panel v). The CST complex structure has been solved by cryo-electron microscopy (Lim et al., 2020), and AF2 correctly predicted the organization of the subcomplex consisting of the CTC1 C-terminal OB-fold, STN1 N-terminal OB-fold, and TEN1 (Fig 1B panel vii), though the position of STN1 C-terminal winged helix-turn-helix domains bound to CTC1 was not predicted.

Shieldin is known to interact with the CST complex that facilitates fill-in of resected DNA by Polymerase α-primase (Mirman et al., 2022a). The AF2 analysis predicted two points of interaction between shieldin and CST: SHLD1-CTC1 (Fig 1B panel vi) and SHLD2-CTC1 (Fig 1B panel v). The predicted SHLD1-CTC1 interaction has been previously described (Mirman et al., 2022a) and disrupting this interface by mutating SHLD1 leucine 20 that is buried within the binding surface ablates this interaction (Mirman et al., 2022b). Our AF2 analysis also predicts an interaction between the third SHLD2 OB-fold with CTC1 that is compatible with SHLD1-SHLD2 binding (Fig 1B panel v), though this region is not sufficient for interaction in yeast-two-hybrid assays (Mirman et al., 2018). As the AF2 protein-protein interaction prediction recapitulated most known interactions involving the shieldin complex, we turned our attention to the entirely novel prediction of the RIF1-SHLD3 binding interface.

### RIF1 HEAT repeats interact with the SHLD3 C-terminal eIF4E-like domain

The AF2 analysis predicted a high confidence interaction between the N-terminal HEAT repeats of RIF1 and the C-terminus of SHLD3. The C-terminal half of SHLD3 is predicted to form a globular domain with structural homology to the translation elongation factor eIF4E (DALI search; Holm and Sander, 1995). All five models scored highly, with pDockQ scores ≥0.5, mean interface PAE ≤11.6 Å, and iPTM scores ≥0.7, a range of values that discriminates between accurate and inaccurate predictions by previous benchmarking studies (Fig 1C; Bryant et al., 2022; Yin et al., 2022). An additional AF2 prediction isolating the SHLD3 C-terminus and the RIF1 residues 1-615 (Fig 2A) further improved scores to pDockQ ≥0.5, mean interface PAE ≤5.8 Å, and iPTM ≥0.91, with high pLDDT scores (parameter reflecting per-residue AF2 prediction confidence; Jumper et al., 2021) throughout and a consistent interface across five separate models (Fig S2A-B). The AF2-predicted interface lies between the first two RIF1 HEAT-repeat helices and the face of the SHLD3 eIF4E-like domain β-sheet with a total buried surface area of 1808 Å^2^ (Fig 2B). The AF2 model of the RIF1 N-terminus and the SHLD3 C-terminus is sterically incompatible with a recently described SHLD3 DNA binding activity (Susvirkar and Faesen, 2022), suggesting that any DNA binding by SHLD3 is mutually exclusive with its RIF1 association (Fig S2C). The RIF1 K315/R316 residues, which are essential for 53BP1 binding (Setiaputra et al., 2022), are distal from the predicted SHLD3 binding interface (Fig S2C).

**Figure 2.**
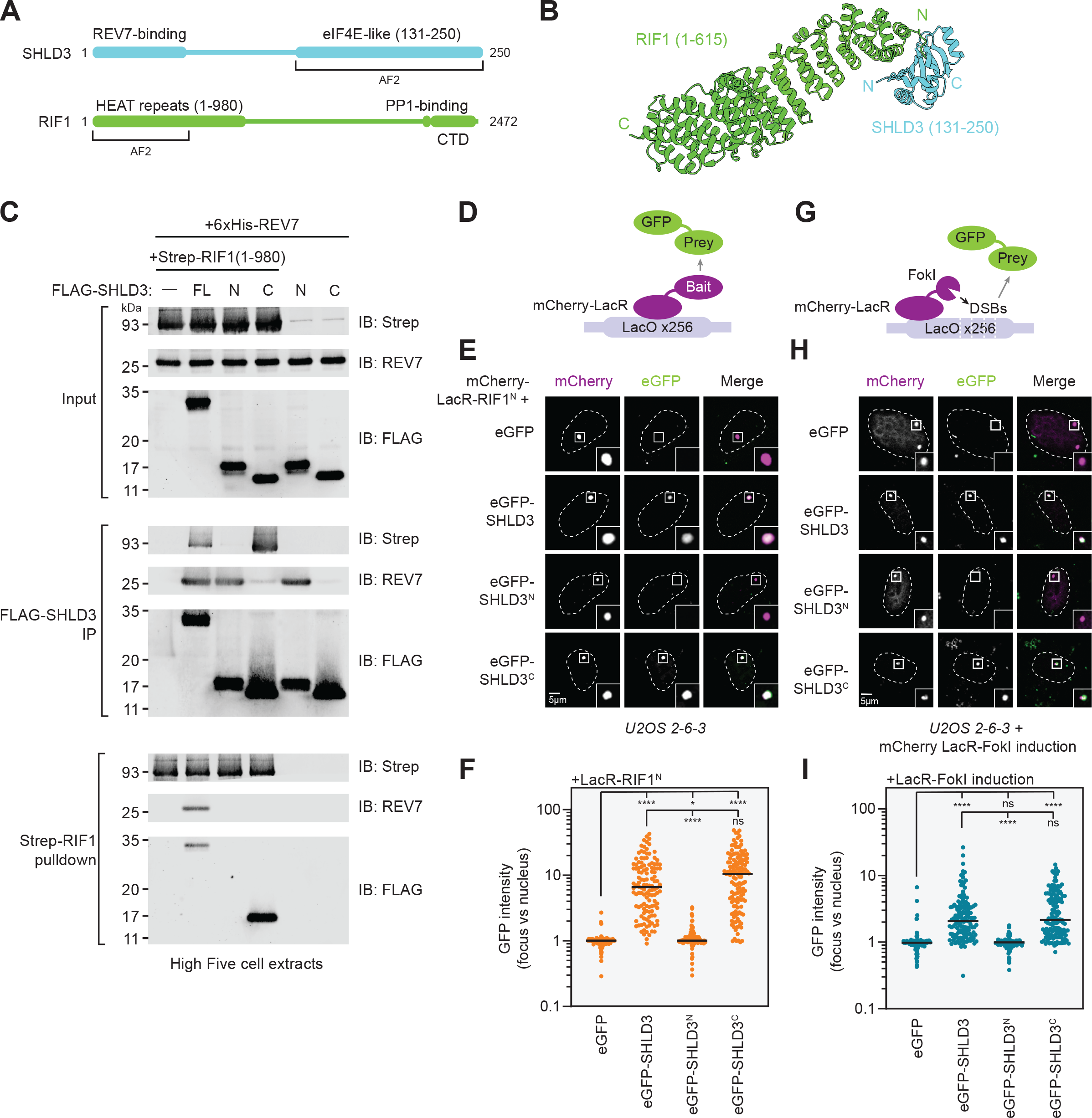
The RIF1 N-terminal HEAT repeats interact with the eIF4E-like C-terminus of SHLD3. A. Schematic of the domain organization of SHLD3 and RIF1, and the fragments used for additional AF2 analysis. B. Top-ranked model predicted by AF2 between RIF1 HEAT repeat residues 1-615 and SHLD3 eIF4E-like domain residues 131-250. See also Fig S2A-C. C. Whole cell extracts of High Five insect cells individually expressing FLAG-SHLD3, Strep-RIF1, and 6xHis-REV7 through baculovirus infection were combined and subjected to FLAG immunoprecipitation or streptactin pulldown and immunoblotted for REV7 or the Strep or FLAG epitopes. Results are representative of three biologically independent experiments. IB: immunoblot, FL: full-length. For SHLD3, N and C correspond to residues 1-125 and 126-250, respectively. D. Schematic of the LacR/LacO assay using plasmid-encoded mCherry-LacR-fused bait and eGFP-fused prey proteins transfected in the U2OS 2-6-3 cell line containing ~256 lac operator (LacO) repeats. E. Representative micrographs of the LacR/LacO assay using mCherry-LacR-RIF1^N^ as bait to evaluate chromatin recruitment of eGFP-tagged SHLD3 variants. SHLD3: residues 2-250. SHLD3^N^: residues 2-125. SHLD3^C^: residues 126-250. RIF1^N^: residues 1-967. See also Fig S2D-F. F. Quantification of E. GFP intensities are presented as a ratio between the average fluorescence intensity within the mCherry-labeled LacR focus and the average nuclear intensity. Bars represent means (*n* = 3 independent experiments with ≥ 39 nuclei imaged each). Analysis was performed by Dunnett’s test comparing against empty vector control. \**\*\*\**: *P* < 0.0001. ns: *P* > 0.05. G. Schematic of the LacR-FokI assay that directs FokI nuclease-mediated DNA double-strand breaks to the U2OS 2-6-3 LacO array to analyze DNA damage localization of eGFP-fused prey proteins encoded by transfected plasmids. H. Representative micrographs of the LacR-FokI assay to evaluate DNA damage recruitment of eGFP-tagged SHLD3 variants after induction of LacR-FokI expression. See also Fig S2G-I. I. Quantification of H. GFP intensities are presented as a ratio between the average fluorescence intensity within the mCherry-labeled LacR-FokI focus and the average nuclear intensity. Bars represent mean (*n* = 3 independent experiments with ≥ 44 nuclei imaged each). Analysis was performed by Dunnett’s test compared to empty vector control. \**\*\*\**: *P* < 0.0001. ns: *P* > 0.05.

To validate the predicted RIF1-SHLD3 interaction *in vitro*, we first used baculovirus-mediated transduction of insect cells to individually express Strep-tagged human RIF1^N^ (residues 1-980), FLAG-tagged SHLD3, and His-tagged REV7 and then performed reciprocal FLAG and Strep pulldowns from combined insect cell extracts to study their interactions. These experiments confirmed previous observations that the N-terminus of SHLD3 (residues 2-125; SHLD3^N^) is necessary and sufficient for REV7 binding, while its eIF4E-like domain-containing C-terminus (residues 126-250; SHLD3^C^) does not participate in this interaction (Fig 2C; Ghezraoui et al., 2018). Consistent with the AF2 prediction, the C-terminus of SHLD3 is sufficient for interacting with the RIF1 N-terminal HEAT-repeat domain encompassed by RIF1^N^. Additionally, RIF1^N^ is unable to bind REV7 in the absence of SHLD3, supporting previous observation that SHLD3 bridges RIF1 and REV7 (Fig 2C; Noordermeer et al., 2018). Together these results show that recombinant SHLD3 interacts with REV7 and RIF1 through its N- and C-terminal domains, respectively.

To determine whether this interaction is recapitulated in cells, we employed the U2OS 2-6-3 cell line that contains an array of ~256 Lac operator (LacO) repeats (Janicki et al., 2004). We expressed the RIF1 HEAT repeats (residues 1-967) fused to mCherry-Lac repressor (LacR) and determined whether it can recruit GFP-labeled SHLD3 to the LacO array (Fig 2D-F, S2D-F). Consistent with the pulldown results, the C-terminus of SHLD3 is necessary and sufficient for its robust recruitment to chromatin by RIF1 (Fig 2E-F). Notably, unlike 53BP1-nucleosome or RIF1-53BP1 interactions, this recruitment occurs without exogenous DNA damage, suggesting that it is independent of DNA damage-induced post-translational modifications.

SHLD3 is responsible for recruiting shieldin to DSB sites (Noordermeer et al., 2018). Since RIF1 is essential for shieldin DNA-damage localization, we hypothesized that the SHLD3 C-terminus is necessary and sufficient for SHLD3 recruitment to DSBs. To test this possibility, we induced DSBs at the LacO array in U2OS 2-6-3 cells by expressing FokI endonuclease fused to mCherry-LacR (Fig 2G, S2G-I; Shanbhag et al., 2010) and determined which region of SHLD3 was essential for its DSB localization. We observed that the SHLD3 C-terminus is essential for its recruitment to sites of FokI-induced DNA breaks (Fig 2H-I, S2G). These observations show that the SHLD3 eIF4E-like domain binds RIF1 HEAT-repeats to recruit shieldin to sites of DNA damage.

### SHLD3 interacts with RIF1 through a highly charged interface

The predicted SHLD3-RIF1 interface contains several conserved basic and acidic residues that are poised to form multiple electrostatic contacts (Fig 3A). We identified five putative electrostatic interactions between conserved pairs of residues (Fig 3A) and expressed SHLD3^C^ variants with alanine substitutions to test the contributions of these predicted electrostatic contacts to the SHLD3-RIF1 interaction. Recombinant SHLD3 variants with W132A, N201A, and D216A mutations were deficient in RIF1^N^ binding in reciprocal pulldown experiments from insect cell extracts, while R166A mutation completely abolished interaction with RIF1^N^ (Fig 3B). We next asked whether these point mutations also disrupt RIF1^N^-SHLD3^C^ interaction in cells. Consistent with the pulldown experiments, the W132A and R166A variants failed to colocalize with LacR-RIF1^N^ at the LacO array, while the N201A and D216A variants were partially defective in colocalization, as shown by weaker intensity foci (Fig 3C, S3A-B). Furthermore, the W132A and R166A variants were unable to accumulate at FokI-induced DSBs, whereas the N201A and D216A variants retained weak localization, suggesting partially impaired interaction with RIF1 (Fig 3D, S3C). In contrast, the S131A substitution did not impact interaction with RIF1^N^ in pulldown assays and this SHLD3 variant was fully proficient in colocalizing with LacR-RIF1^N^ at the LacO array as well as being recruited to FokI-induced DSBs. Transient expression of these SHLD3 variants did not affect the localization of endogenous RIF1 to FokI-induced DSBs (Fig S3D-E), consistent with shieldin being recruited downstream of RIF1. In summary, we evaluated the AF2 prediction that five SHLD3 residues facilitate RIF1-SHLD3 interaction through both *in vitro* pulldown and cellular recruitment assays and determined that R166 is essential for RIF1 binding, W132, N201, and D216 are important, while S131 is dispensable for this activity.

**Figure 3.**
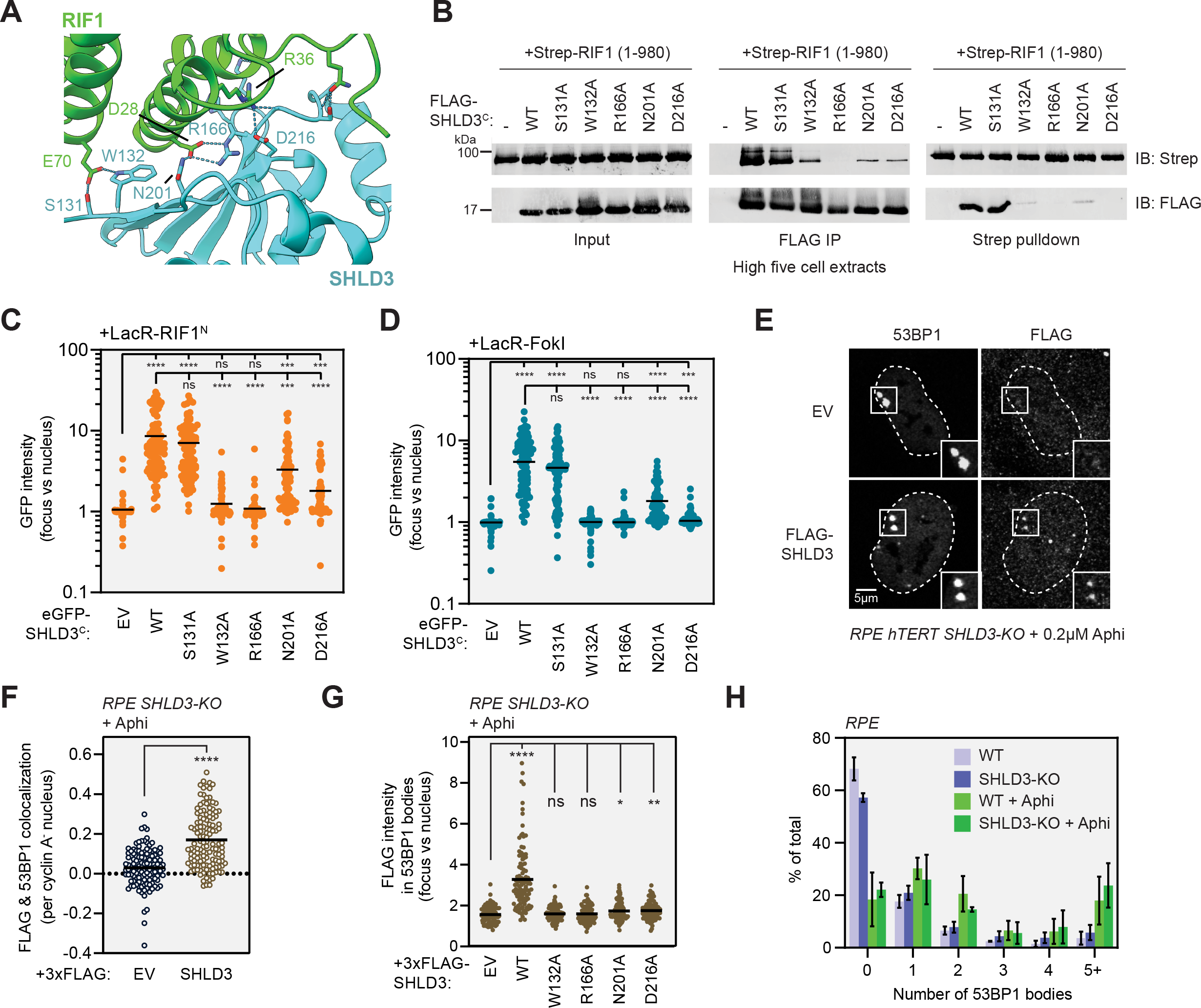
SHLD3 binds RIF1 through polar interactions that are essential for recruitment to DNA breaks and 53BP1 bodies. A. Close-up view of the hydrogen bond and salt bridge network between conserved RIF1 and SHLD3 residues in the AF2-predicted model. Electrostatic interactions are depicted as dashed cyan lines. B. Whole cell extracts of High Five insect cells individually expressing the wild-type (WT) or indicated alanine substitution FLAG-SHLD3^C^ (residues 126-250) variants and Strep-RIF1^N^ (residues 1-980) through baculovirus infection were combined and subjected to FLAG immunoprecipitation or streptactin pulldown and immunoblotted for the Strep or FLAG epitopes. The results are representative of two independent experiments. IB: immunoblot. IP: immunoprecipitation. C. Quantification of LacR/LacO assay measuring recruitment of eGFP-SHLD3^C^ wild-type (WT) or alanine substitution variants to LacO arrays by mCherry-LacR-RIF1^N^. GFP intensities are presented as a ratio between the average fluorescence intensity within the mCherry-labeled LacR-RIF1^N^ focus and the average nuclear intensity. Bars represent means (*n* = 3 independent experiments with ≥19 nuclei imaged each). See also Fig S3A-B. EV: empty vector. Unless otherwise indicated, all statistical analysis in this figure was performed using Dunnett’s test. \**\*\*\**: *P* < 0.0001. ***: *P* < 0.001. ns: *P* > 0.05. D. Quantification of LacR-FokI assay measuring recruitment of eGFP-SHLD3^C^ wild-type (WT) or alanine substitution variants to sites of DNA double-strand breaks induced by mCherry-LacR-FokI. GFP intensities are presented as a ratio between the average fluorescence intensity within the mCherry-labeled LacR-FokI focus and the average nuclear intensity. Bars represent means (*n* = 3 independent experiments with ≥ 39 nuclei imaged each). See also Fig S3C-E. EV: empty vector. E. Representative micrographs of immunofluorescence experiments (from 3 biologically independent replicates) analyzing localization of SHLD3 to 53BP1 bodies. RPE-hTERT p53-KO SHLD3-KO cells were complemented by transduction of lentivirus encoding FLAG-tagged SHLD3. The cells were then treated with 200 nM aphidicolin (Aphi) for 24 h and processed for immunofluorescence microscopy using antibodies against 53BP1, FLAG, and cyclin A. 53BP1 bodies are defined as distinct foci visible in cyclin A-negative cells. EV: empty vector. F. Quantification of FLAG and 53BP1 colocalization from E performed in CellProfiler. Pearson’s correlation coefficients (PCC) were calculated for pixels within each cyclin A-negative, 53BP1 body-positive nucleus between the 53BP1 and FLAG channels. Each point represents the PCC value of an individual nucleus. Bars represent means. Three biologically independent replicates with ≥ 30 nuclei imaged each were performed. Analysis was performed using Welch’s t-test. ****: *P* < 0.0001. EV: empty vector. G. Quantification of immunofluorescence experiments measuring FLAG intensities within 53BP1 bodies in RPE-hTERT p53-KO SHLD3-KO cells expressing lentivirus-encoded FLAG-SHLD3 wild-type (WT) or alanine substitution variants and treated with 200 nM Aphi for 24 h. Individual points represent the ratio between average FLAG intensity within 53BP1 bodies and the average nuclear intensity. Bars represent means. Three biologically independent replicates with ≥ 30 nuclei imaged each were performed. Analysis was performed using Dunnett’s test compared to empty vector (EV) control. ****: *P* < 0.001, **: *P* = 0.009, *: *P* = 0.03, ns: *P* > 0.05. See also Fig S3F-G. H. Quantification of cells containing the indicated number of 53BP1 bodies in RPE-hTERT p53-KO WT and SHLD3-KO cells with or without 24-hour 200 nM aphidicolin (Aphi) treatment. Bars represent means ± s.d. (*n* = 3 biologically independent replicates with ≥ 30 nuclei imaged each).

### SHLD3 is recruited to 53BP1 bodies through its interaction with RIF1

In addition to focal accumulation around DSBs, 53BP1 marks DNA lesions in G1 associated with replication stress in the preceding cell cycle (Lukas et al., 2011). These 53BP1 bodies are large and their formation requires many of the same DNA damage signaling factors involved in 53BP1 localization to DSBs such as the kinase ATM, histone H2AX phosphorylation, and MDC1 (Harrigan et al., 2011). 53BP1 bodies arise from mitotic passage of under-replicated DNA at difficult-to-replicate loci such as common fragile sites (Harrigan et al., 2011). A recent study implicated RIF1 and shieldin in the prevention of aberrant HR at 53BP1 bodies, leading to delayed resolution of these lesions (Spies et al., 2019). However, the presence of shieldin at such bodies has not been described. We generated *SHLD3* knockouts (*SHLD3-KO*) in the RPE1 hTERT p53^−/−^ background and restored SHLD3 expression using a lentivirus encoding 3xFLAG-tagged SHLD3 (FLAG-SHLD3; Fig S3F). We induced mild DNA replication stress with a low dose (0.2 μM) of the B-family DNA polymerase inhibitor aphidicolin (Wright and Brown, 1990) and analyzed SHLD3 localization into 53BP1 bodies by immunofluorescence using Pearson Correlation Coefficient measurements between pixels within each nucleus (Fig 3E-F). FLAG-SHLD3 shows robust colocalization with 53BP1 bodies. Measurements of FLAG-SHLD3 intensity within 53BP1 bodies found that SHLD3 variants defective in RIF1 binding were deficient in 53BP1 colocalization (Fig 3G, S3G). Notably, the N201A and D216A variants that retained partial RIF1 binding show very slight but statistically significant recruitment to 53BP1 bodies (EV mean = 1.56 vs N201A, D216A means = 1.74, 1.76; *P* = 0.03, 0.01, respectively). Since RIF1 promotes formation of 53BP1 bodies in response to aphidicolin-induced replication stress (Watts et al., 2020), we investigated whether loss of SHLD3 affects 53BP1 body formation. In both the absence and presence of aphidicolin, *SHLD3-KO* cells do not display significant differences in the number of 53BP1 bodies (Fig 3H), consistent with SHLD3 recruitment being downstream of 53BP1. These observations suggest that similar mechanisms underlie shieldin recruitment to both 53BP1 bodies and DSB sites, and that the importance of RIF1 for 53BP1 body formation is independent of shieldin accumulation at these sites.

### Dissecting the RIF1 surface that facilitates SHLD3 binding

Next, we sought to validate the region of RIF1 that interacts with SHLD3. AF2 analysis predicts that the two N-terminal α-helices of the HEAT-repeat domain participate in SHLD3 interactions (Fig 4A). Consistent with this prediction, LacR fused to a truncated form of the RIF1 HEAT-repeat domain containing only the N-terminal 567 residues was fully proficient in recruiting full-length SHLD3 to LacO arrays (Fig 4A, S4A-B). However, deletion of the first 173 residues from this protein (yielding RIF1 174-567), which includes the predicted SHLD3-binding region, abolished its ability to interact with SHLD3 as determined by their lack of cellular colocalization at the LacO array (Fig 4A, S4A-B). To further interrogate the predicted RIF1-SHLD3 binding interface, we expressed LacR-RIF1^N^ proteins containing alanine substitutions at residues E70, D28, and R36 that are predicted to form hydrogen bonds with SHLD3 residues W132, R166 + N201, and D216, respectively (Fig 3A). The LacR-RIF1^N^ D28A and R36A variants were highly defective in recruiting SHLD3 to LacO arrays (Fig 4B, S4C-D). Unexpectedly, LacR-RIF1 E70A was fully proficient in interacting with SHLD3 despite the importance of SHLD3 W132 in both LacO and DSB recruitment (Fig 3C-D). This observation indicates that the SHLD3 W132 residue facilitates RIF1-SHLD3 binding through interactions other than the predicted E70-W132 hydrogen bond. We then individually expressed SHLD3^C^ with the D28A and R36A RIF1^N^ variants in insect cells and tested their ability to interact through reciprocal pulldown experiments in combined cell extracts. Consistent with the LacR experiments, both D28A and R36A RIF1^N^ variants were deficient in copurifying with SHLD3^C^ in vitro, with D28A having the most severe defect (Fig 4C). These results point to the predicted RIF1^D28^-SHLD3^R166^ electrostatic interaction as an essential component of the binding interface, and indicate that the predicted RIF1^R36^-SHLD3^D216^ interaction also likely contributes to the SHLD3-RIF1 interaction.

**Figure 4.**
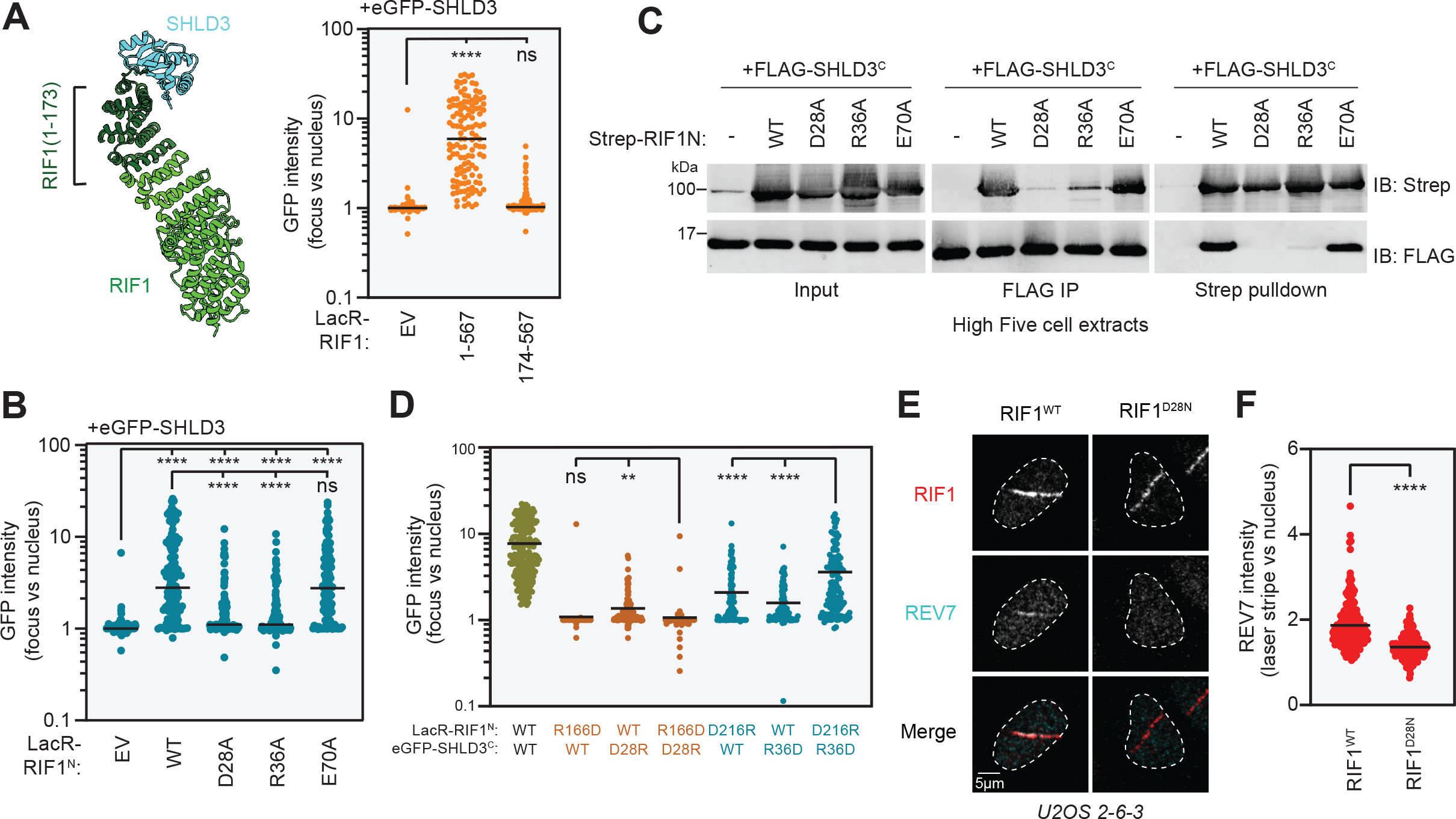
RIF1 binds SHLD3 through polar residues within its extreme N-terminus. A. Left: Representation of the region of the RIF1 N-terminal HEAT repeats that is predicted to bind the SHLD3 eIF4E-like domain. Residues 1-173 are highlighted. Right: Quantification of LacR/LacO assay measuring eGFP-SHLD3 recruitment to LacO arrays by the indicated truncated mCherry-LacR-RIF1^N^ variants. GFP intensities are presented as a ratio between the average fluorescence intensity within the mCherry-labeled LacR-RIF1^N^ focus and the average nuclear intensity. Bars represent means (*n* = 3 independent experiments with ≥ 37 nuclei imaged each). Analysis was performed using Dunnett’s test compared to empty vector (EV) control. ****: *P* < 0.001, ns: *P* = 0.54. See also Fig S4A-B. EV: empty vector. B. Quantification of LacR/LacO assay measuring eGFP-SHLD3 recruitment to LacO arrays by mCherry-LacR-RIF1^N^ (residues 1-967) or the indicated alanine substitution variants. GFP intensities are presented as a ratio between the average fluorescence intensity within the mCherry-labeled LacR focus and the average nuclear intensity. Bars represent means (*n* = 3 independent experiments with ≥ 34 nuclei imaged each). Analysis was performed using Dunnett’s test compared to empty vector (EV) and wild-type RIF1^N^ (WT) control. ****: *P* < 0.001, ns: *P* = 0.47. See also Fig S4C-D. EV: empty vector. C. Whole cell extracts of High Five insect cells individually expressing the FLAG-SHLD3^C^ (residues 126-250) and the indicated alanine substitution Strep-RIF1 (residues 1-980) variants through baculovirus infection were combined and subjected to FLAG immunoprecipitation or streptactin pulldown and immunoblotted for the Strep or FLAG epitopes. The results are representative of two independent experiments. IB: immunoblot. IP: immunoprecipitation. D. Quantification of LacR/LacO assay measuring eGFP-SHLD3^C^ recruitment to LacO arrays by mCherry-LacR-RIF1^N^ with both transfected plasmids bearing charge-reversal mutations. Bars represent means (*n* = 3 independent experiments with ≥ 39 nuclei imaged each). Analysis was performed using Dunnett’s multiple comparisons test. ****: *P* < 0.001, **: *P* = 0.045, ns: *P* = 0.98. See also Fig S4E-F. E. Representative micrographs of UV laser microirradiation experiments measuring DNA damage recruitment of REV7. DNA damage was induced in U2OS 2-6-3 cells through irradiation in the form of linear stripes and analyzed by immunofluorescence microscopy with RIF1 and REV7 antibodies. See also Fig S4G-H. F. Quantification of E. REV7 immunofluorescence intensities are presented as a ratio between the average fluorescence intensity within the RIF1-labeled irradiation stripe and the average nuclear intensity. Only RIF1-positive nuclei are quantified. Bars represent means (*n* = 2 independent experiments with ≥ 90 nuclei imaged each). Analysis was performed using Welch’s t-test. ****: *P* < 0.0001.

We next probed the RIF1^D28^-SHLD3^R166^ and RIF1^R36^-SHLD3^D216^ electrostatic interaction pairs through charge-reversal mutation using the LacO/LacR chromatin localization assay (Fig 4D, S4E-F). As predicted, individual charge-reversal mutations within the two electrostatic interaction pairs result in defective mCherry-LacR-colocalizing eGFP-SHLD3^C^ focus formation. Combining the RIF1^N^ R166D and SHLD3^C^ D28R mutations did not rescue SHLD3^C^ focus formation, suggesting either that the combined mutations did not restore the electrostatic interaction, or that other essential interactions are associated with these residues. Indeed, RIF1^D28^ is also predicted to form a hydrogen bond with SHLD3^N201^ (Fig 3A-B). Strikingly, combining the RIF1^N^ R36D and SHLD3^C^ D216R variants partially rescues SHLD3^C^ focus formation, providing strong evidence that this salt bridge was accurately predicted by AF2 and that it plays a key role in the RIF1-SHLD3 interaction.

Thus far the experiments we performed utilized cellular models overexpressing RIF1 and SHLD3 variants. To ensure that our observations apply to endogenously produced proteins, we used a dCas9-guided base editor (Koblan et al., 2018) to generate U2OS 2-6-3 cells carrying homozygous D28N mutations at the endogenous *RIF1* locus (Fig S4G). The D28N mutation did not alter RIF1 expression (Fig S4H). Since the isolated homozygous RIF1-D28N clones lost the LacO array, we monitored REV7 recruitment to UV laser microirradiation sites instead of at FokI-induced DSBs (Fig 4E) and found that RIF1^D28N^ failed to recruit REV7 to sites of laser microirradiation (Fig 4E-F). We conclude that shieldin recruitment to DSB sites depends on the physical and direct interaction between SHLD3 and RIF1 modeled by AF2.

### SHLD3-RIF1 interaction is essential for class switch recombination

We next investigated whether mutations abolishing SHLD3-RIF1 binding abrogates shieldin function. We evaluated antibody class switch recombination (CSR) in CH12F3 mouse B cell lymphoma cells. Upon stimulation with a cocktail of anti-CD40 antibody, TGF-β, and IL-4, CH12F3 cells rapidly undergo CSR to convert its expressed immunoglobulin isotype from IgM to IgA (Nakamura et al., 2006). CSR involves induction of DSBs at switch regions within the immunoglobulin heavy chain locus and ligation of distal breaks leading to exon recombination (Methot and Di Noia, 2017). This process is reliant on the 53BP1-RIF1-shieldin pathway, whose disruption leads to excessive resection of CSR-induced DSBs into the coding regions of the immunoglobulin locus (Ghezraoui et al., 2018; Ling et al., 2020). Accordingly, CRISPR-Cas9-generated CH12F3 *Shld3^−/−^* cells are severely deficient in CSR (Fig S5A-B; Noordermeer et al., 2018).

We evaluated the ability of SHLD3 variants deficient in RIF1 interaction to undergo CSR (Fig 5A-C, S5C-D). CH12F3 *Shld3^−/−^* cells transduced with lentivirus expressing wild-type SHLD3 regained CSR activity. Consistent with DSB localization and *in vitro* pulldown results, the R166A variant that is completely unable to interact with RIF1 (Fig 3B-C) did not support CSR. Interestingly, the W132A, N201A, and D216A variants that retained partial interaction with RIF1 (Fig 3B-C) showed differing CSR activity. While the W132A variant did not rescue IgA class switching, both N201A and D216A variants showed comparable CSR to the wild-type complementation (Fig 5B-C). Our results suggest that the RIF1-SHLD3 interaction is essential for shieldin-dependent CSR, and possibly shieldin function, downstream of 53BP1.

**Figure 5.**
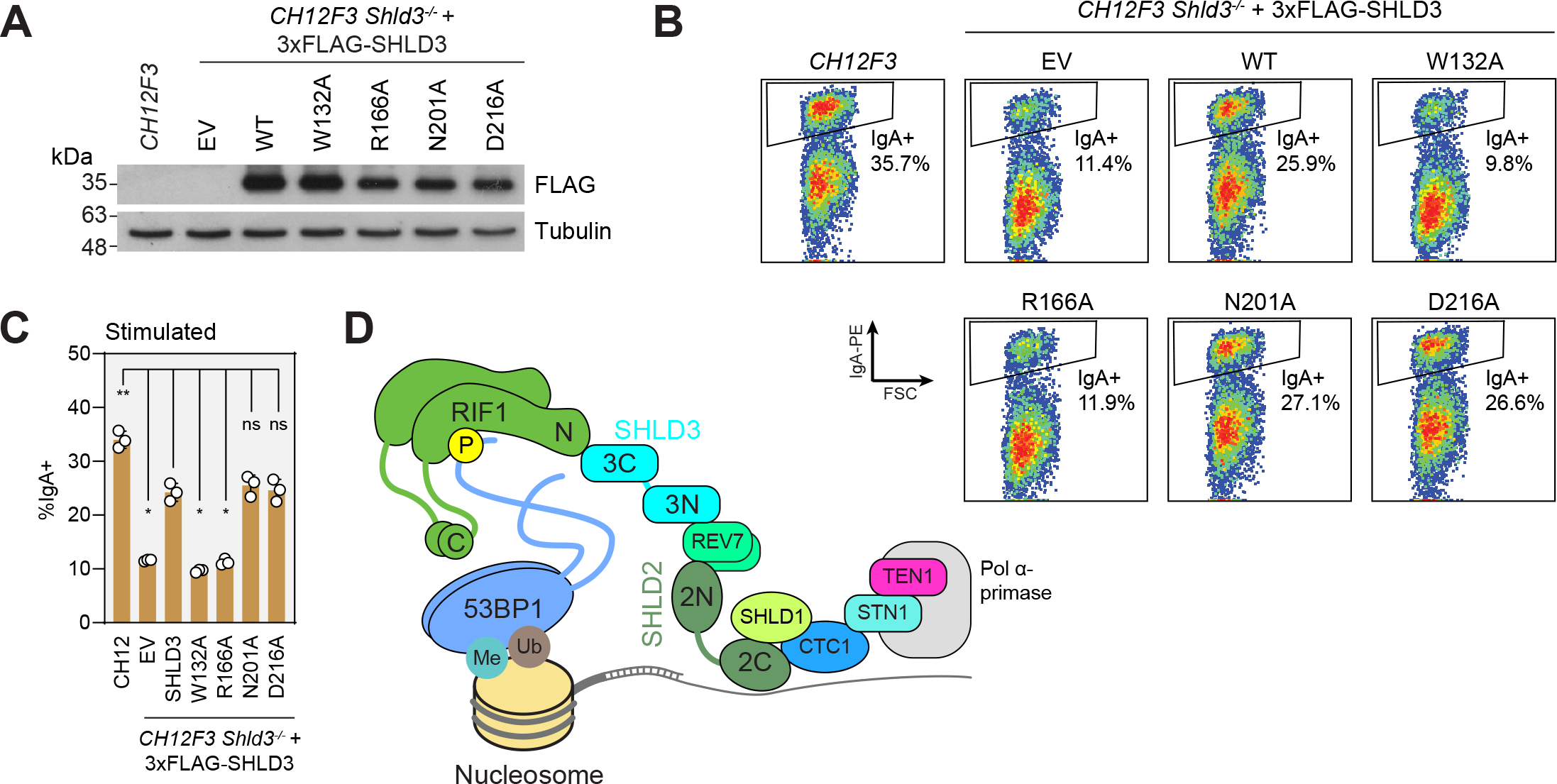
RIF1-SHLD3 interaction is essential for shieldin function in CSR. A. Immunoblot of whole cell extracts of CH12F3 *Shld3^−/−^* cells stably transduced with lentivirus encoding the indicated FLAG-tagged SHLD3 alanine substitution variants. Lysates were probed for FLAG and tubulin (loading control). Representative of two immunoblots. EV: empty vector. WT: wild-type. B. Representative flow cytometry density plots (from three biologically independent experiments) measuring IgA expression in stimulated CH12F3 *Shld3^−/−^* cells expressing FLAG-tagged SHLD3 alanine substitution variants. Values shown are %IgA^+^ cells. WT: wild-type. EV: empty vector. FSC: forward scatter. See also Fig S5C-D. C. Quantification of class switch recombination data shown in B. Bars represent means ± s.d., *n* = 3 biologically independent experiments. Analysis was performed using Dunnett’s test compared to *Shld3^−/−^* cells complemented with wild-type SHLD3. **: *P* < 0.01, *: *P* < 0.05, ns: *P* > 0.05. EV: empty vector. WT: wild-type. D. Schematic of the molecular basis for 53BP1-RIF1-shieldin-CST recruitment to sites of DNA damage.

### Molecular determinants of 53BP1-RIF1-shieldin-CST recruitment to DNA damage sites

In this study we used an exhaustive pairwise matrix of AlphaFold2-Multimer predictions to augment our knowledge of the protein-protein interactions that define the 53BP1-RIF1-shieldin-CST pathway. These predictions provide the structural basis for 53BP1 phosphorylation-dependent recognition by RIF1 (Setiaputra et al., 2022), 53BP1 dimerization through its oligomerization domain (Zgheib et al., 2009), RIF1 dimerization through its C-terminus (Moriyama et al., 2018), and SHLD1 binding to CTC1 (Mirman et al., 2022b). We interrogated the novel prediction of the RIF1-SHLD3 binding interface through multiple experiments that all proved to be consistent with the AF2 model. Furthermore, we determined that the physical interaction between RIF1 and SHLD3 is essential for 53BP1 pathway function in CSR. These findings enhance our understanding of the molecular basis for 53BP1-RIF1-shieldin-CST assembly with putative structural information that shows remarkable agreement with experimental data (Fig 5D) and represent yet another validation of the potential of AF2 to discover novel biologically relevant protein-protein interactions.

## Experimental Methods

### Cell lines

U2OS 2-6-3 cells (Shanbhag et al., 2010) were cultured in McCoy’s 5A (Modified) Medium (Gibco) supplemented with 10% fetal bovine serum (FBS; Wisent) and 50 IU/mL penicillin, 50 μg/mL streptomycin (Wisent). CH12F3-2 cells (referred to as CH12F3) were cultured in RPMI 1640 (Gibco) supplemented with 10% FBS, 50 IU/mL penicillin, 50 μg/mL streptomycin, 5% NCTC-109 (Gibco), 60 μM β-mercaptoethanol (Sigma-Aldrich). CH12F3-2 *Shld3^−/−^* Clone 2 was previously generated (Noordermeer et al., 2018). RPE1 hTERT p53-KO FLAG-Cas9 and 293T cells were cultured in Dulbecco’s Modified Eagle Medium (DMEM; Gibco) supplemented with 10% FBS, 50 IU/mL penicillin, 50ug/ml streptomycin, 1x GlutaMax (Gibco), 1xMEM non-essential amino acids (MEM-NEAA; Gibco). Sf9 and High Five insect cells were maintained in suspension in I-Max (Wisent).

RPE1 hTERT p53-KO SHLD3-KO FLAG-Cas9 cells were generated by transfecting RPE1 hTERT p53-KO FLAG-Cas9 cells with *in vitro*-transcribed sgRNA using Lipofectamine RNAiMAX (Thermo-Fisher) according to manufacturer’s instructions using 0.6 μg each of two sgRNAs (sgSHLD3 #1: GGTGATCTTTTAGGTCTGAG, sgSHLD3 #2: TGAATTGTAGCATTACAAGA). sgRNAs were generated by *in vitro* transcription using TranscriptAid T7 High Yield Transcription Kit (Thermo Scientific) and cleaned using the Agencourt RNAClean XP Kit (Beckman-Coulter) according to manufacturer’s instructions. After transfection, individual clones were collected and knockouts were confirmed by PCR amplification and the ICE CRISPR analysis tool (Synthego).

U2OS 2-6-3 D28N were generated by electroporation of pCMV_BE4max (Addgene #112093; Koblan et al., 2018) and phU6-gRNA expression cassette encoding the guide sequence AAGCGTCAGTCTGCCCTCCA (Addgene #53188; Kabadi et al., 2014). After 3 days of recovery, cells were plated at low density to isolate colonies. Editing was analyzed by PCR amplification of the edited region and Sanger sequencing.

### Plasmids

eGFP-SHLD3 truncation-expressing plasmids were generated using Gibson cloning to delete fragments from pcDNA5-FRT/TO-eGFP-SHLD3 (Noordermeer et al., 2018). Amino acid substitutions in the C-terminal (126-250) truncation were created via QuikChange site-directed mutagenesis (Agilent). pDEST-mCherry-LacR-RIF1 (1-567) truncations were generated by deletion PCR of the pDEST-mCherry-LacR-RIF1 (1-967) plasmid. Amino acid substitutions in pDEST-mCherry-LacR-RIF1 (1-967) were created using QuikChange site-directed mutagenesis.

pAC8-FLAG-SHLD3 truncation plasmids were generated using Gibson cloning (New England Biolabs) and a pAC8-FLAG-SHLD3 template (Setiaputra et al., 2022). Amino acid substitutions of pAC8-FLAG-SHLD3 were made via site-directed mutagenesis. Amino acid substitutions of Strep-RIF1 (1-980) were generated using site-directed mutagenesis on a pFastBac-Strep-RIF1(1-980) template (Setiaputra et al., 2022).

The pHIV-3xFLAG-NAT plasmid was generated by inserting a 3xFLAG N-terminal tag into the pHIV-NAT-hCD52 plasmid (Willis et al., 2018) by ligation of annealed oligos into the NotI and XmaI sites. The SHLD3 (2-250) coding sequence was ligated into the NheI and XmaI sites of pHIV-3xFLAG-NAT.

### AlphaFold2-Multimer pairwise matrix screen for protein-protein interactions

Amino acid sequences for human 53BP1, RIF1, SHLD1, SHLD2, SHLD3, REV7, CTC1, STN1, TEN1, and ASTE1 were retrieved from UniProt. The default sequences were used except for SHLD2, where the longer isoform (Q86V20-2) was used. Longer proteins were manually split—avoiding cutting at structured sites—to accommodate graphical memory limitations. 53BP1 was divided into 4 fragments (1-600, 601-1200, 1201-1715, 1716-1927), RIF1 into 4 fragments (1-570, 571-1200, 1200-1800, 1800-2472), SHLD2 into 2 fragments (1-420, 421-904), and CTC1 into 2 fragments (1-546, 547-1217). Joint FASTA files for every unique pair (including self-pairs) was generated using a Python script and used as input files for AlphaFold2-Multimer (AF2) prediction.

The LocalColabFold v1.4 implementation of AF2 was used (Mirdita et al., 2022) on a cloud 8xTesla V100 16GB GPU instance from Lambda Labs, using the following command: colabfold_batch –num-recycle 3 –num-models 5 –model-type AlphaFold2-multimer-v2 <fasta input folder>/ <output folder>/

The confidence of each predicted interface was analyzed by a Python script measuring pDockQ (Bryant et al., 2022) and mean interface predicted aligned error (PAE). Mean interface predicted aligned error was determined by identifying every pair of residues whose Cβ atoms (or Cα if glycine) were within 9 Å (identification of interface residues was adapted from pDockQ.py; https://gitlab.com/ElofssonLab/FoldDock/-/tree/main/src from Bryant et al., 2022), then calculating the average PAE value across all residue pairs. Individual models that have pDockQ scores >0.23 and mean interface PAE scores <15 Å were classified as potential interactors. Unique protein pairs with at least 4/5 models meeting this cutoff or having previous supporting experimental evidence were classified as high-confidence interactors. Protein structures were visualized with UCSF ChimeraX (Goddard et al., 2018). Modeling phosphate groups, H-bond prediction, surface hydrophobicity calculation, and multi-model alignments were performed using built-in ChimeraX functions. Additional individual AF2 predictions were performed using the ColabFold Google Colab AlphaFold2_mmseqs2 (https://colab.research.google.com/github/sokrypton/ColabFold/blob/main/AlphaFold2.ipynb) sheet.

### Lentivirus generation and infection

Lentiviruses were generated in 293T cells by co-transfecting the pHIV-3xFLAG-NAT viral targeting vector with plasmids encoding RRE, REV, and VSV-G using the TransIT-LT1 Transfection Reagent (Mirus). Viral supernatants were collected, filtered through a 0.45 μm syringe filter, and snap frozen in liquid nitrogen. Viral infections were performed in the presence of 8 μg/mL polybrene (Sigma-Aldrich) for 24 h, after which the media was refreshed and selection with 0.2 μg/mL nourseothricin (Gold Biotechnology) was performed until uninfected controls no longer survived, upon which selection was no longer maintained.

### SHLD3-RIF1 co-immunoprecipitations

Baculoviruses were generated in *Spodoptera frugiperda* Sf9 cells (Thermo Fisher) using the Bac-to-Bac method for the pFastBac-derived vectors or, for the pAC8-derived vectors, cotransfecting them with linearized viral DNA (Abdulrahman et al., 2009). The proteins were expressed in *Trichoplusia ni* High Five cells (Expression Systems) by baculoviral expression. Individual High Five cell cultures were infected so that they would express a single recombinant protein.

36 h post-infection the cells were harvested by centrifugation. The cells were resuspended in lysis buffer (50 mM Tris-HCl pH 8, 200 mM NaCl, 1 mM PMSF, 1 mM TCEP, 1x SIGMAFAST protease inhibitor tablet) and lysed by sonication prior to being centrifuged at high speed (21,000 xg, 30 min, 4°C) and the supernatant collected. Lysate containing SHLD3 recombinant protein was mixed with lysate containing RIF1 recombinant protein, REV7 recombinant protein, or both. The lysate mixtures were applied to FLAG M2 beads (Sigma-Aldrich) or Strep-Tactin Superflow high capacity resin (IBA Lifesciences) and incubated with rotation at 4°C for 1 h. The resin was washed twice, each wash including a 20 min incubation with rotation at 4°C. The pulldowns were eluted from the FLAG and Strep resins using 0.1 mg/mL 3xFLAG peptide (GlpBio) or 2.5 mM desthiobiotin (Sigma-Aldrich), respectively. The results were analysed by immunoblotting.

### U2OS-2-6-3 SHLD3-RIF1 colocalization assay

2.5-3.0 x 10^5^ U2OS-2-6-3 cells containing a LacO array of 256 repeated LacO sequences (Shanbhag et al., 2010) were plated on glass coverslips in 6-well plates. The next day, the cells were transfected as follows: 1 μg eGFP-SHLD3 constructs, 1 μg mCherry-LacR-RIF1 constructs and 6 μL Lipofectamine 2000 (Thermo Scientific) were incubated in Opti-MEM (Gibco) for 20 min at room temperature before being added to the cells whose media had been changed to McCoy’s with 10% FBS and no antibiotics. Two hours later, the transfection media was replaced by fresh McCoy’s with 10% FBS and no antibiotics. Two days after transfection, the cells were washed in PBS, then incubated for 10 min on ice in 1 mL nuclear pre-extraction buffer (20 mM HEPES pH 7.4, 20 mM NaCl, 5 mM MgCl_2_, 0.5% NP-40, 1 mM DTT, and 1x cOmplete EDTA-free protease inhibitor cocktail [Roche]), and washed again in PBS. The coverslips were then incubated for 10 min at room temperature in 1 mL 4% formaldehyde (Thermo Scientific) in PBS and washed 3 times in PBS.

Images for this and other microscopy experiments were acquired on a Zeiss LSM780 laser scanning confocal microscope on either a 40x or 63x Plan-Apochromat objective lens (specific to the experiment). Images were quantified using ImageJ to determine the average fluorescence intensities of the nuclear region colocalizing with the mCherry-LacR signal and the average fluorescence intensity within the nuclear boundary. 1 was added to both values before calculating the ratio to minimize confounding effects of dividing with extremely low intensity values.

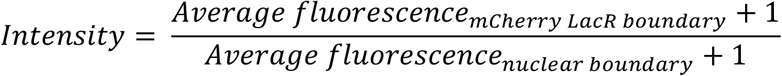

Any adjustments to image brightness or contrast were applied to the entire image, and to the same extent between all images of the same experiment.

### U2OS-2-6-3 FokI focus recruitment assay

2.5 x 10^5^ U2OS-2-6-3 cells containing a LacO array of 256 repeated LacO sequences and an inducible mCherry-LacR-FokI (Shanbhag et al., 2010) were plated on glass coverslips in 6-well plates. The next day, the cells were transfected as follows: 2 μg eGFP-SHLD3 constructs and 6 μl Lipofectamine 2000 (Thermo Scientific) were incubated in Opti-MEM for 20 min at room temperature before being added to the cells whose media had been changed to McCoy’s with 10% FBS and no antibiotics. Two hours later, the media was changed to fresh McCoy’s with 10% FBS and no antibiotics. Two days after transfection, the expression of mCherry-LacR-FokI was induced by the addition of 10 μg/mL 4-hydroxytamoxifen (Sigma-Aldrich) and 1 μM of Shield-1 peptide (Clontech, Mountain View CA) for 4 h. Cells were washed in PBS, then incubated for 10 min on ice in 1 mL nuclear pre-extraction buffer (20 mM HEPES pH 7.4, 20 mM NaCl, 5 mM MgCl_2_, 0.5% NP-40, 1 mM DTT, and 1x cOmplete EDTA-free protease inhibitor cocktail [Roche]), and washed again in PBS. The coverslips were then incubated for 10 min at room temperature in 1 mL 4% formaldehyde in PBS and washed 3 times in PBS. Images were acquired and analyzed as described above.

### Immunofluorescence

Fixed cells cultured on glass coverslips were placed in a humidified chamber and incubated in blocking solution (PBS + 0.2% cold water fish gelatin + 0.5% BSA) for 30 min. The blocking solution was replaced with primary antibody diluted in blocking solution and incubated for 2 h at room temperature. The coverslips were washed three times for a total of 15 minutes in PBS then incubated for 1 h at room temperature with secondary antibody diluted in blocking solution. The coverslips were washed three times for a total of 15 minutes in PBS then mounted onto glass slides using ProLong Gold Antifade mounting media with DAPI (Invitrogen). Coverslips were imaged as described above.

### Class switch recombination assay

Class switch recombination assays were performed essentially as previously described (Noordermeer et al. 2018). 1 x 10^5^ CH12F3-2 cells were plated in 24-well plates in growth medium supplemented with 1 μg/mL anti-CD40 antibody (eBioscience), 1 ng/ml TGF-β (R&D Systems), and 10 ng/mL mIL-4 (R&D Systems). After 48 h, cells were harvested, stained with anti-IgA-PE and fixed with 4% formaldehyde. Fluorescence signal was acquired on an Attune NxT Flow Cytometer (Thermo-Fisher). Data was analyzed using FlowJo software (BD Biosciences).

### Analyzing localization to 53BP1 bodies

RPE cells were grown on glass coverslips and treated with 200 nM aphidicolin (Sigma-Aldrich) for 24 h. Coverslips were then washed with PBS, fixed with 4% formaldehyde for 10 min at room temperature, washed again 3x with PBS, permeabilized with 0.3% Triton X-100 in PBS for 30 min at room temperature, and washed 3x with PBS. Coverslips were stained with antibodies to FLAG, 53BP1, and cyclin A. Cyclin A-positive cells were discarded from the analysis. For Pearson Correlation Coefficient calculation, the MeasureColocalization functionality of CellProfiler v4.2.5 (Stirling et al., 2021) was used within each masked nucleus to calculate the pixel correlation between the 53BP1 and FLAG channels. For FLAG intensity in 53BP1 bodies measurement, 53BP1 foci in cyclin A-negative nuclei were manually masked in ImageJ, and the mean FLAG fluorescence intensity within the mask was divided by the mean nuclear FLAG fluorescence intensity. Number of 53BP1 bodies were counted automatically by CellProfiler.

### Laser microirradiation

Laser microirradiation experiments were performed as previously described (Setiaputra et al., 2022) except that coverslips were harvested 1 h after irradiation.

**Table 1.** Antibodies used in this study.

| Reagent or resource | Source | Identifier | Antibody concentration |
| --- | --- | --- | --- |
| Mouse monoclonal anti-FLAG M2 | Sigma-Aldrich | Cat #: F1804;<br>RRID:AB_262044 | IB: 1:500 |
| Mouse monoclonal anti-FLAG M2-HRP | Sigma-Aldrich | Cat #: A8592 | IB: 1:1000 |
| Rabbit polyclonal anti-Strep Tag II | Abcam | Cat #: ab76949;<br>RRID:AB_1524455 | IB: 1:1000 |
| Rabbit recombinant monoclonal anti-MAD2L2/REV7 | Abcam | Cat #: ab180579;<br>RRID:AB_2890174 | IF: 1:500 |
| Goat polyclonal anti-eGFP | Gift from L. Pelletier | N/A | IB: 1:5000 |
| Rabbit polyclonal anti-mCherry | Abcam | Cat #: ab167453;<br>RRID:AB_2571870 | IB: 1:1000 |
| Mouse monoclonal anti-RIF1 (B-3) | Santa Cruz Biotechnology | Cat #: Sc-515573 | IF: 1:100 |
| Rabbit polyclonal anti-RIF1 | Bethyl Laboratories | Cat #: A300-569A | IB: 1:1000 |
| Rabbit polyclonal anti-Phospho-Histone H2A.X (ser139) | Cell Signaling Technologies | Cat #: 2577L | IF: 1:500 |
| Mouse monoclonal anti-Cyclin A | Santa Cruz Biotechnology | Cat #: Sc-271682 | IF: 1:200 |
| Rabbit polyclonal anti-53BP1 | Bethyl Laboratories | Cat #: A300-272A | IF: 1:1000 |
| Goat anti-FLAG | Abcam | Cat #: ab1257 | IF: 1:3000 |
| Goat anti-mouse IgA-PE | Southern Biotech | Cat #: #1040-09 | FC: 1:100 |
| Mouse monoclonal anti-α-tubulin (DM1A) | Cell Signaling Technologies | Cat #: 3873S | IB: 1:2000 |
| Alexa Fluor 647 goat anti-mouse | Invitrogen | Cat #: A21236 | IF: 1:1000 |
| Alexa Fluor 647 goat anti-rabbit | Invitrogen | Cat #: A21244 | IF: 1:1000 |
| IRDye 680RD-Goat polyclonal anti-mouse IgG | LI-COR Biosciences | Cat #: 926-68070 | IB: 1:10000 |
| IRDye 800CW-Goat polyclonal anti-rabbit IgG | LI-COR Biosciences | Cat #: 926-32211 | IB: 1:10000 |
| IRDye 800CW-Donkey polyclonal anti-rabbit IgG | LI-COR Biosciences | Cat #: 926-32213 | IB: 1:10000 |
| IRDye 800CW-Donkey polyclonal anti-goat IgG | LI-COR Biosciences | Cat #: 925-32214 | IB: 1:10000 |
| IRDye 680RD-Donkey polyclonal anti-mouse IgG | LI-COR Biosciences | Cat #: 926-68072 | IB: 1:10000 |
| IRDye 680RD-Donkey polyclonal anti-rabbit IgG | LI-COR Biosciences | Cat#: 926-68073 | IB: 1:10000 |

## Acknowledgments

We thank R. Szilard for critical reading of this manuscript and other members of the Durocher lab for valuable discussion. We thank R. Greenberg for the U2OS 2-6-3 cell line and R. Scully for the pHIV-NAT-hCD52 plasmid. D.S. was funded by a Cancer Research Society Next Generation Scholarship for most of this work. D.D. is a Canada Research Chair (Tier 1) and work in the D.D. lab was funded from grants from the Canadian Institutes for Health Research (CIHR, PJT-180438) and the Krembil Foundation to D.D..

## Supplementary figure legends

**Figure S1.**
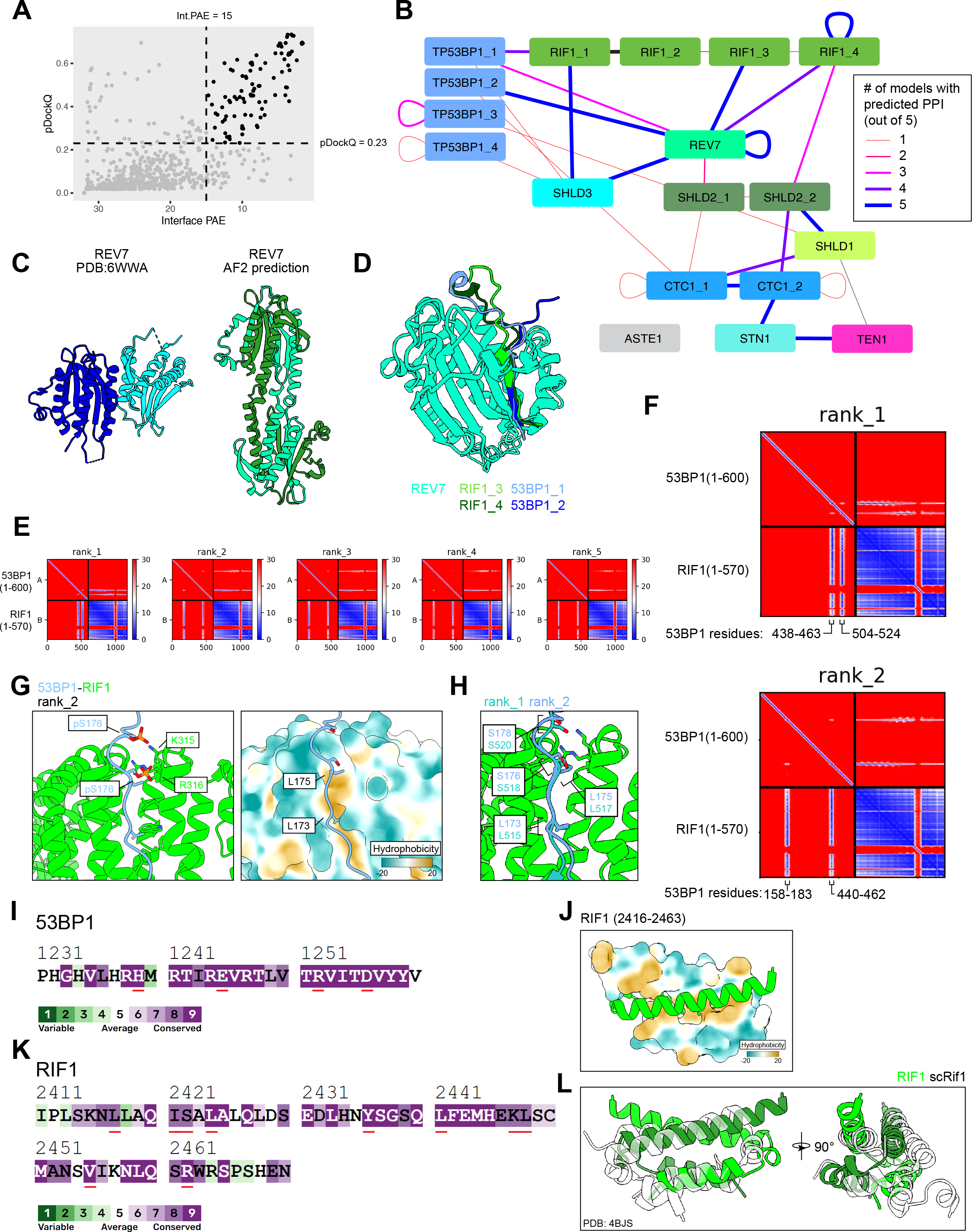
Data supporting the AlphaFold2-Multimer pairwise matrix screen for protein-protein interactions in the 53BP1-RIF1-shieldin-CST pathway. A. Scatter plot of pDockQ and interface predicted aligned error (PAE) scores of individual models from the interaction prediction screen. B. Schematic of predictions that meet the cutoff scores of pDockQ >0.23 and interface PAE <15 Å. Nodes are proteins and fragments used for each prediction. Edges link the two chains used for each prediction. Edge thickness and color correspond to the number of predictions (out of 5 for each pair) meeting the score cutoffs. C. Comparison of REV7 dimer structures determined from X-ray crystallography (Left, PDB ID: 6WWA) and from AlphaFold2 prediction (right). D. Superimposition of predicted heterodimeric structures between REV7 and RIF1 fragments 3 and 4 and 53BP1 fragments 1 and 2. E. PAE plots of the predicted 53BP1 (fragment 1; 1-600) and RIF1 (fragment 1; 1-570), ranked by predicted template model (pTM) scores. F. Close-ups of two PAE plots from E, with the 53BP1 regions of high PAE confidence labeled. G. Predicted structures of the 53BP1-RIF1 interface. Left, phosphate groups modeled onto two serines of 53BP1 whose phosphorylation are known to be essential for interaction with RIF1. Right, surface representation of RIF1 colored by hydrophobicity. H. Superimposition of the 53BP1-RIF1 interface from two different predicted models. I. ConSurf sequence conservation analysis of the indicated 53BP1 region. Residues important for the predicted 53BP1 oligomerization are underlined in red. J. Predicted structure of the RIF1 dimerization interface, one monomer is shown with surface representation colored by hydrophobicity. K. ConSurf sequence conservation analysis of the indicated RIF1 region. Residues important for the predicted RIF1 oligomerization are underlined in red. L. Superimposition of the RIF1 oligomerization domain and the experimentally-determined structure (translucent) of the corresponding region in *S. cerevisiae* Rif1 (PDB ID: 4BJS).

**Figure S2.**
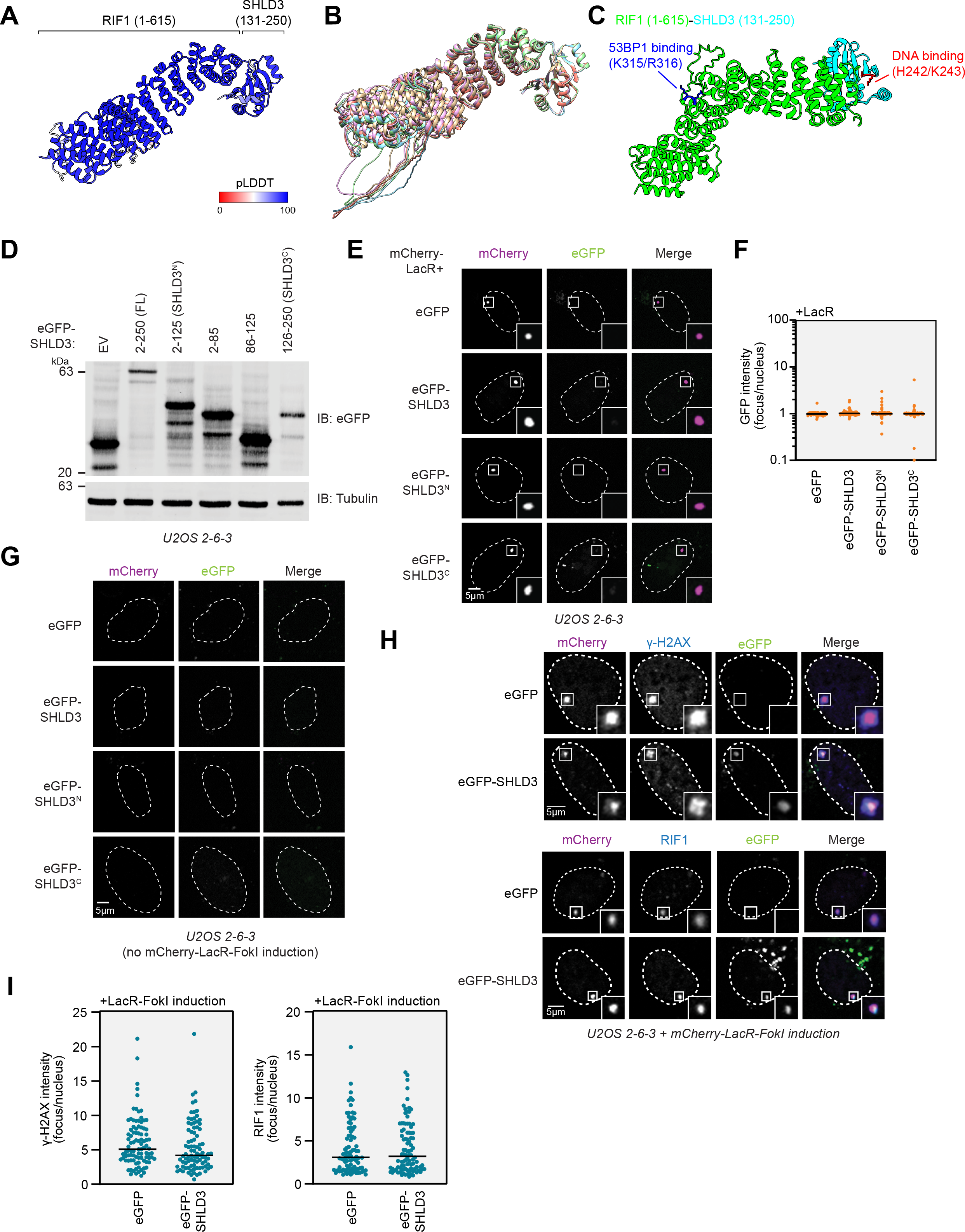
Data supporting the SHLD3 C-terminus as necessary and sufficient for RIF1 binding and recruitment to sites of DNA damage. A. Top-ranking model of RIF1 (1-615) and SHLD3 (131-250) colored by pLDDT score. B. Five AF2-predicted models of RIF1 (1-615) and SHLD3 (131-250) superimposed, aligned by the SHLD3 chain. C. Top-ranking AF2 model of RIF1 (1-615) and SHLD3 (131-250), highlighting residues associated with SHLD3 DNA binding and RIF1 53BP1 phosphopeptide binding. D. Immunoblot of whole cell extracts of U2OS 2-6-3 cells transfected with plasmids encoding the indicated eGFP-tagged SHLD3. Lysates were probed for eGFP and tubulin (loading control). Representative of two biologically independent immunoblots. E. Representative micrographs (from three biologically independent experiments) of the LacR/LacO assay using mCherry-LacR as bait to evaluate chromatin recruitment of eGFP-tagged SHLD3 variants as a control for the experiments using mCherry-LacR-RIF1^N^ in Fig 2E-F. F. Quantification of E. GFP intensities are presented as a ratio between the average fluorescence intensity within the mCherry-labeled LacR focus and the average nuclear intensity. Bars represent mean (*n* = 3 independent experiments with ≥ 39 nuclei imaged each). G. Representative micrographs of control experiments for the LacR-FokI assay in Fig 2H-I to evaluate DNA damage recruitment of eGFP-tagged SHLD3 variants. LacR-FokI expression was not induced, and no mCherry-LacR-FokI or eGFP-SHLD3 foci were detected. H. Representative micrographs of the LacR-FokI assay to evaluate DNA damage induction after mCherry-LacR-FokI expression. U2OS 2-6-3 cells were transfected with plasmids encoding eGFP-SHLD3 and treated with 4-hydroxytamoxifen and Shield-1 peptide to induce mCherry-LacR-FokI expression. The cells were then analyzed for γ-H2AX focus formation colocalizing with mCherry-LacR-FokI as a proxy for DNA double-strand break formation through immunofluorescence (top). Colocalization of mCherry-LacR-FokI focus with endogenous RIF1 was also assessed (bottom). I. Quantification of H. Immunofluorescence intensities are presented as a ratio between the average fluorescence intensity within the mCherry-labeled LacR focus and the average nuclear intensity. Bars represent mean (*n* = 2 independent experiments with ≥ 40 nuclei imaged each).

**Fig S3.**
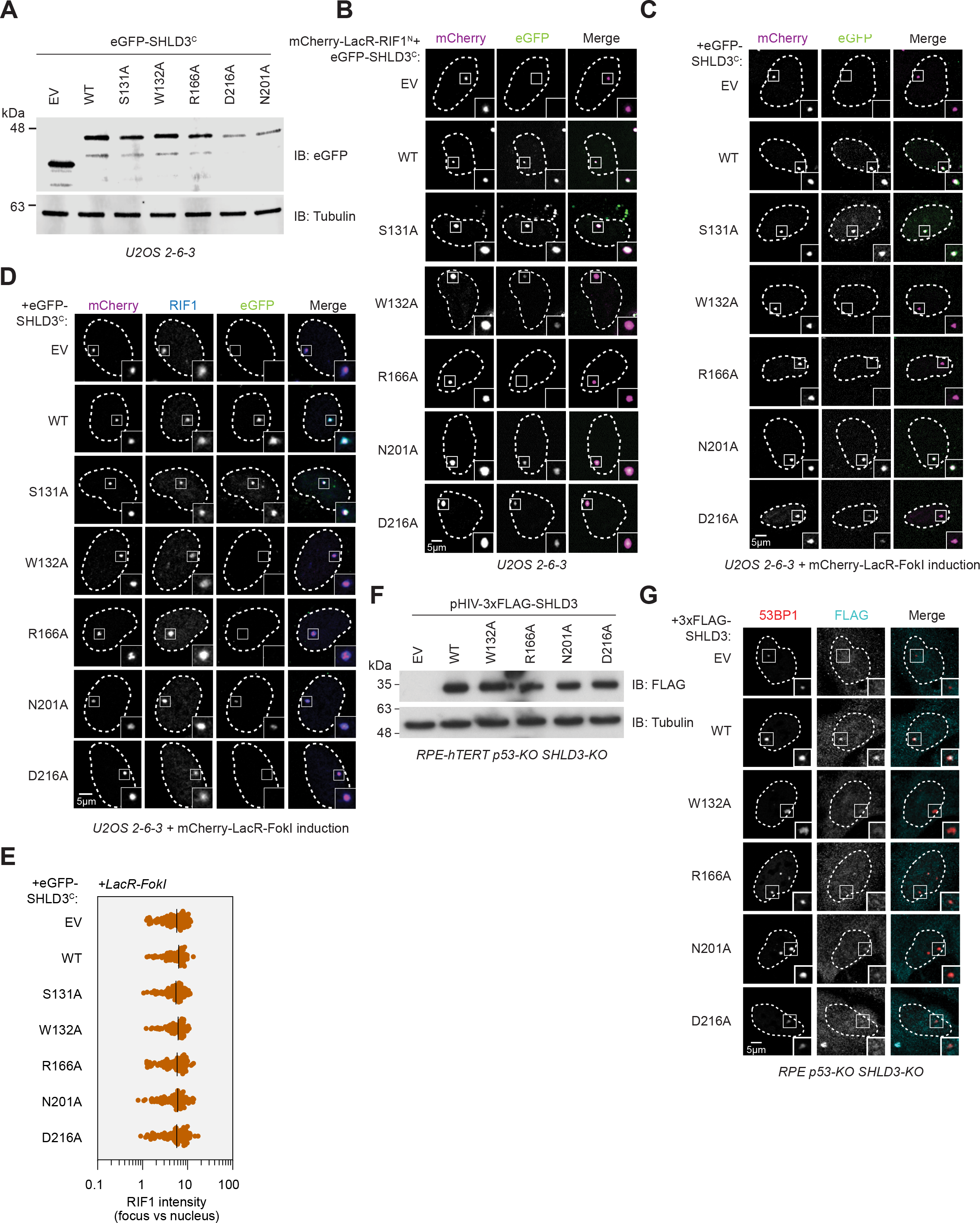
Data supporting the polar interactions that are essential for SHLD3-RIF1 binding. A. Immunoblot of whole cell extracts of U2OS 2-6-3 cells transfected with plasmids encoding the indicated eGFP-tagged SHLD3^C^ (residues 126-250). Lysates were probed for eGFP and tubulin (loading control). Representative of two immunoblots. EV: empty vector. WT: wild-type. B. Representative micrographs (of three biologically independent experiments) of the LacR/LacO assay using mCherry-LacR-RIF1^N^ as bait to evaluate chromatin recruitment of eGFP-tagged SHLD3^C^ alanine substitution variants shown in Fig 3C. SHLD3^C^: residues 126-250. RIF1^N^: residues 1-967. EV: empty vector. WT: wild-type. C. Representative micrographs (of three biologically independent experiments) of the LacR-FokI assay to evaluate DNA damage recruitment of eGFP-tagged SHLD3^C^ alanine substitution variants after induction of LacR-FokI expression shown in Fig 3D. SHLD3^C^: residues 126-250, EV: empty vector. WT: wild-type. D. Representative micrographs (of two biologically independent experiments) of the LacR-FokI assay to evaluate DNA damage recruitment of endogenous RIF1 after induction of LacR-FokI expression in the presence of exogenously expressed eGFP-tagged SHLD3^C^ alanine substitution variants. SHLD3^C^: residues 126-250, EV: empty vector. WT: wild-type. E. Quantification of D. RIF1 immunofluorescence intensities are presented as a ratio between the average fluorescence intensity within the mCherry-labeled LacR focus and the average nuclear intensity. Bars represent means (*n* = 2 independent experiments with ≥ 41 nuclei imaged each). F. Immunoblot of whole cell extracts of RPE-hTERT p53-KO SHLD3-KO cells stably transduced with lentivirus encoding the indicated 3xFLAG-tagged SHLD3 alanine substitution variants. Lysates were probed for FLAG and tubulin (loading control). Representative of two immunoblots. EV: empty vector. WT: wild-type. G. Representative micrographs of immunofluorescence experiments (from 3 biologically independent replicates) analyzing localization of FLAG-SHLD3 alanine substitution variants stably expressed in RPE-hTERT p53-KO SHLD3-KO cells by lentivirus transduction. Cells were treated with 200 nM aphidicolin for 24 hours and processed for immunofluorescence microscopy using antibodies raised against 53BP1, FLAG, and cyclin A. 53BP1 bodies are defined as distinct foci visible in cyclin A-negative cells. Quantification is shown in Fig 3G.

**Fig S4.**
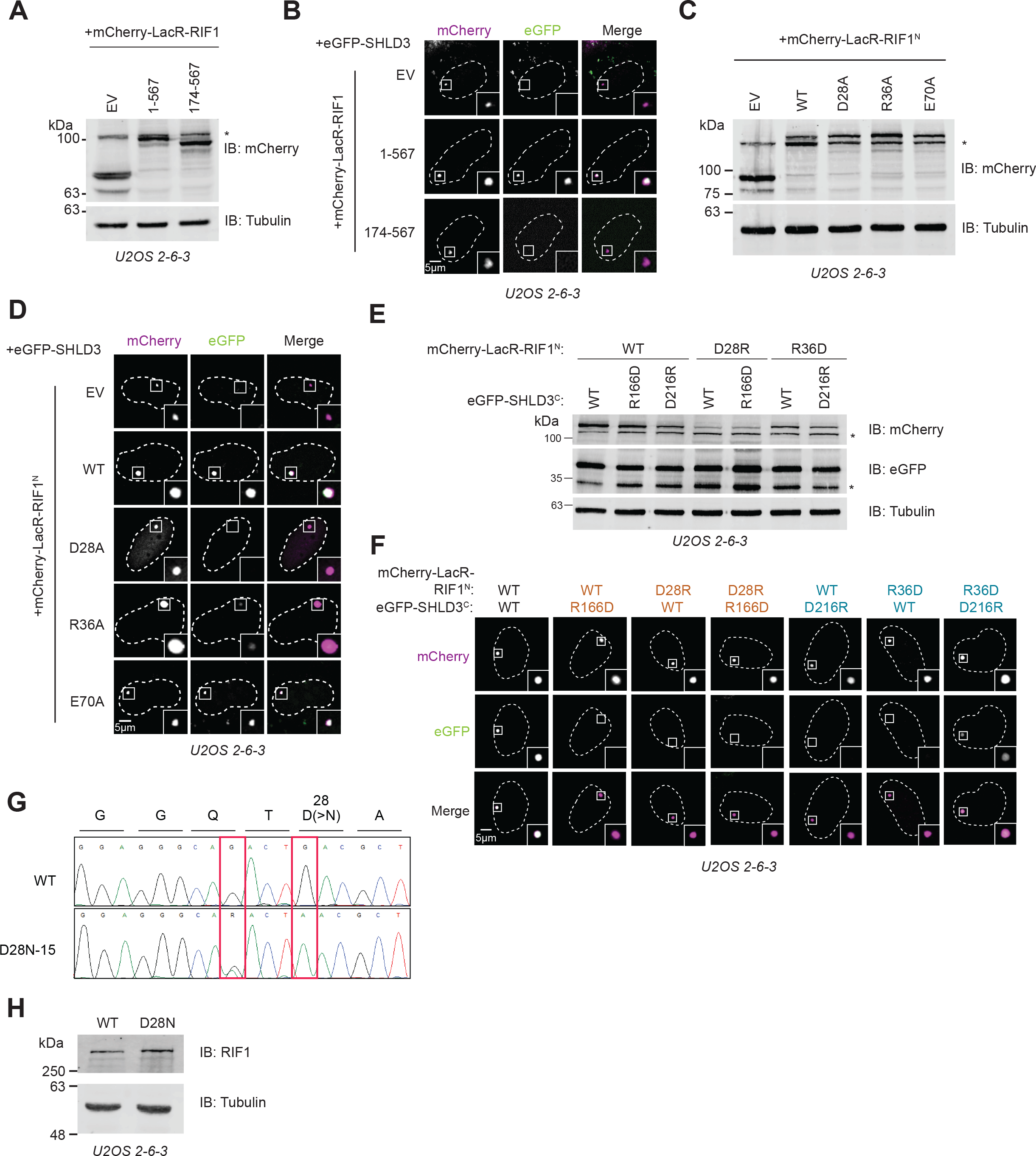
Data supporting the extreme N-terminus of RIF1 binding SHLD3 through polar residues. A. Immunoblot of whole cell extracts of U2OS 2-6-3 cells transfected with plasmids encoding the indicated mCherry-LacR-fused RIF1 truncations. Lysates were probed for mCherry and tubulin (loading control). IB: immunoblot, EV: empty vector. *: nonspecific band. B. Representative micrographs (of three biologically independent experiments) of the LacR/LacO assay using the indicated mCherry-LacR-fused RIF1 variants to evaluate their ability to recruit eGFP-SHLD3 to chromatin shown in Fig 4A. EV: empty vector. C. Immunoblot of whole cell extracts of U2OS 2-6-3 cells transfected with plasmids encoding the indicated mCherry-LacR-fused RIF1^N^ (residues 1-967) alanine variants. Lysates were probed for mCherry and tubulin (loading control). Representative of two immunoblots. IB: immunoblot, EV: empty vector. *: nonspecific band. D. Representative micrographs (of three biologically independent experiments) of the LacR/LacO assay using the indicated mCherry-LacR-fused RIF1^N^ variants to evaluate their ability to recruit eGFP-SHLD3 to chromatin shown in Fig 4B. EV: empty vector. E. Immunoblot of whole cell extracts of U2OS 2-6-3 cells transfected with plasmids encoding the indicated mCherry-LacR-RIF1^N^ and eGFP-SHLD3 variants. Lysates were probed for mCherry, eGFP, and tubulin (loading control). IB: immunoblot. EV: empty vector. WT: wild-type. *: nonspecific band. F. Representative micrographs (of three biologically independent experiments) of the LacR/LacO assay using the indicated mCherry-LacR-RIF1^N^ and eGFP-SHLD3^C^ variants to evaluate their colocalization at LacO arrays shown in Fig 4D. EV: empty vector. WT: wild-type. G. Sanger sequencing chromatograms of PCR products amplified from U2OS 2-6-3 cells with and without subjecting it to base-editing to introduce endogenous D28N mutations. A single clone was isolated that contained the desired mutation (D28N-15). Red boxes highlight induced mutations. The first heterozygous G>A mutation is silent. The second homozygous G>A mutation results in the desired D28N substitution. WT: wild-type. H. Immunoblot of whole cell extracts of U2OS 2-6-3 cells with or without base editing to introduce endogenous D28N mutation. Lysates were probed for RIF1 and tubulin (loading control). Representative of two immunoblots. IB: immunoblot.

**Fig S5.**
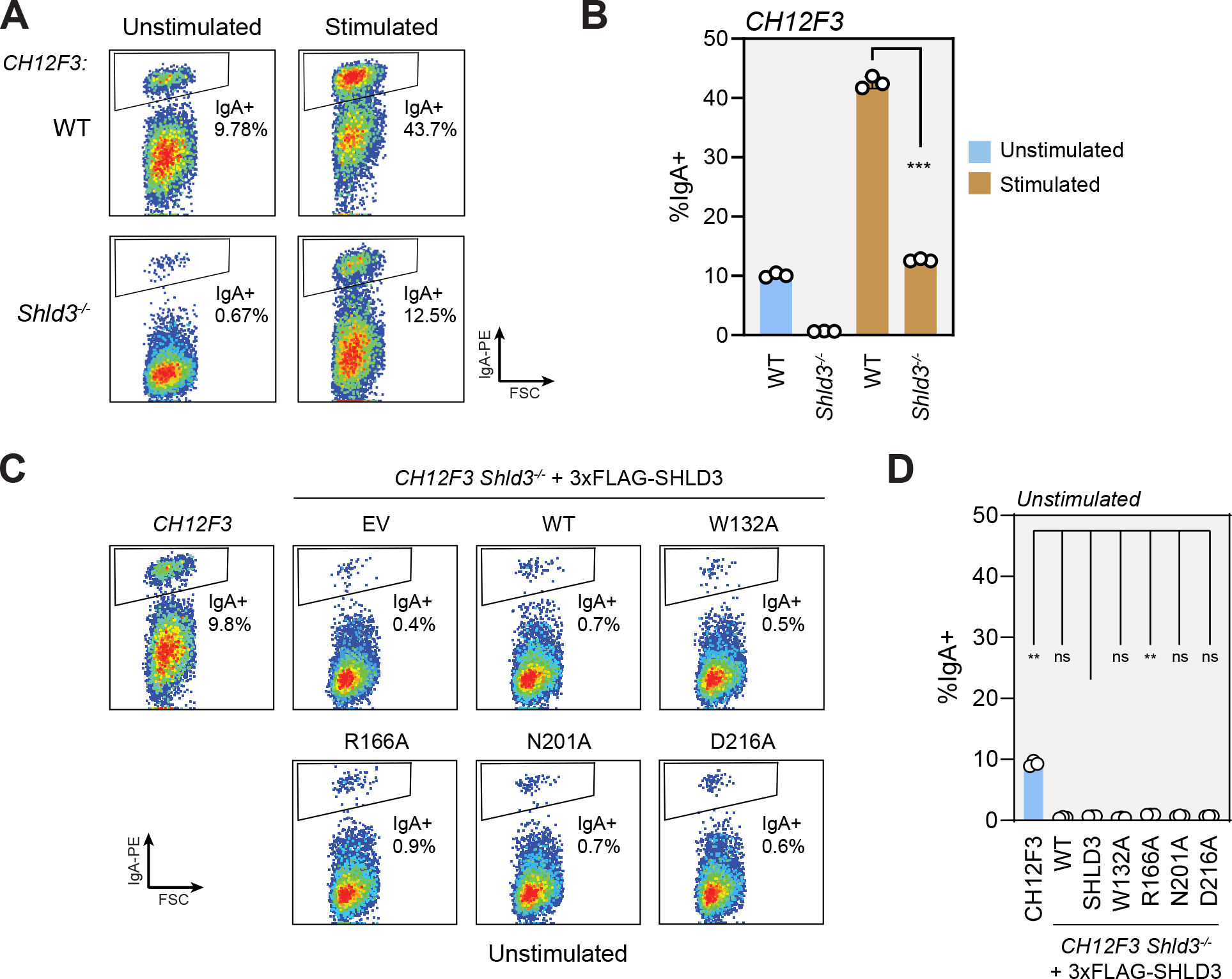
Data supporting the importance of SHLD3-RIF1 binding for class switch recombination. A. Representative flow cytometry density plots (from three biologically independent experiments) measuring IgA expression in unstimulated and stimulated CH12F3 wild-type or *Shld3^−/−^* cells. Values shown are %IgA^+^ cells. EV: empty vector. FSC: forward scatter. WT: wild-type. B. Quantification of class switch recombination data shown in A. Bars represent means ± s.d., *n* = 3 biologically independent experiments. Analysis was performed using Welch’s two-tailed t test. ***: *P* = 0.0002. C. Representative flow cytometry density plots (from three biologically independent experiments) measuring IgA expression in unstimulated CH12F3 wild-type or *Shld3^−/−^* cells stably transduced with lentivirus encoding FLAG-tagged SHLD3 alanine substitution variants. Control experiment for Fig 5B. Values shown are %IgA^+^ cells. EV: empty vector. WT: wild-type. D. Quantification of class switch recombination data shown in C. Bars represent means ± s.d., *n* = 3 biologically independent experiments. Analysis was performed using Dunnett’s multiple test compared against *Shld3^−/−^* cells complemented with wild-type SHLD3. **: *P* < 0.01, ns: *P* > 0.05. EV: empty vector. WT: wild-type.

